# Chromatin topology control by a muscle-specific ribosomal protein

**DOI:** 10.64898/2026.06.23.733628

**Authors:** Makoto Nakamura, Xiaoxin Chen, Shun Yao, Lynette Xin Chan, Ruan Hongmei, Alexis Boulinguiez, Nav Lally, Hao Wu, Kara Kodani, Kentaro Hirose, James Pirruccello, Alberto Malerba, Yifan Cheng, Vasanth Vedantham, Longzhi Tan, Jeffrey E. Olgin, Di Lang, Guo N. Huang

## Abstract

Three-dimensional genome organization stabilizes cell-type-specific gene expression, yet the tissue-restricted factors that maintain chromatin insulation remain poorly understood. Here, we identify the muscle-specific ribosomal protein Rpl3l as an unexpected nuclear regulator of genome architecture in atrial cardiomyocytes. Rpl3l is enriched in the nucleus and nucleolus, where it binds its own genomic locus and stabilizes a CTCF-anchored chromatin boundary that represses the T-type calcium channel gene *Cacna1h*. Loss of *Rpl3l* weakens local chromatin insulation, increases long-range contacts across the *Rpl3l*–*Cacna1h* locus, derepresses *Cacna1h*, and increases susceptibility to atrial fibrillation (AF), which is suppressed by pharmacological inhibition of T-type calcium channels. Furthermore, AF-associated RPL3L variants exhibit impaired nucleolar localization, reduced rRNA binding, and defective repression of *CACNA1H* in human iPSC-derived atrial cardiomyocytes. Together, these findings reveal a ribosomal protein–chromatin axis linking genome insulation to ion-channel dosage control and cardiac rhythm stability, expanding the repertoire of cell-type-specific genome architecture regulators.

**Highlights:** The muscle-specific ribosomal protein Rpl3l exhibits unexpected nuclear and nucleolar enrichment.

Rpl3l stabilizes a CTCF-anchored chromatin boundary to maintain atrial-specific local genome insulation.

Loss of Rpl3l derepresses *Cacna1h* which encodes a T-type Ca2+ channel and increases atrial fibrillation susceptibility.

Atrial fibrillation-associated RPL3L variants impair nucleolar targeting and *CACNA1H* repression

## Introduction

Cell identity depends not only on transcription factors and epigenetic states, but also on the three-dimensional (3D) organization of the genome (1, 2). Chromatin is partitioned into topologically associating domains (TADs) and insulated neighborhoods that constrain regulatory communication and help prevent inappropriate gene activation (3, 4). Although core architectural systems such as CTCF are broadly expressed (5), boundary strength and long-range contacts vary across lineages and change during differentiation (6, 7). How differentiated tissues stabilize locus-selective chromatin insulation to maintain precise gene dosage remains unclear.

The prevailing framework posits that lineage-determining transcription factors, epigenetic modifiers, and chromatin remodelers sculpt cell-type-specific chromatin landscapes by modulating accessibility and enhancer activity (1, 2). Within this view, three-dimensional genome organization is shaped by factors that directly bind DNA or modify chromatin. Consistent with this framework, lineage-determining factors such as MyoD, Pax5, SATB2, and TBX5 have been implicated in higher-order genome organization in skeletal muscle, B cells, neurons, and cardiomyocytes, respectively (8, 9, 10, 11). Together, these studies support a model in which cell-type-specific genome architecture is established and maintained by proteins that directly regulate transcription or chromatin state. However, whether lineage-specific genome architecture can also be reinforced by molecules outside these canonical regulatory categories remains largely unexplored. In particular, it remains unknown whether proteins traditionally assigned to other cellular compartments can exert noncanonical nuclear functions that directly influence higher-order chromatin structure.

Here we identify the muscle-enriched ribosomal paralog Rpl3l as a noncanonical nuclear regulator of chromatin topology in atrial cardiomyocytes. Although best known for roles in ribosome biology and translation, Rpl3l localizes to the nucleus/nucleolus and stabilizes a CTCF-anchored chromatin boundary across the *Rpl3l*–*Cacna1h* region, thereby maintaining repression of the T-type calcium channel gene *Cacna1h*. Loss of *Rpl3l* weakens local insulation, increases contacts across the boundary, derepresses *Cacna1h*, enhances T-type Ca^2+^ current, and drives atrium-selective electrophysiological remodeling and atrial fibrillation (AF) susceptibility. Moreover, AF-associated RPL3L variants exhibit impaired nucleolar localization and reduced rRNA binding and fail to repress *CACNA1H*, linking subnuclear targeting to channel control.

Together, these findings uncover a ribosomal protein–based mechanism that enforces locus-selective 3D genome insulation to safeguard atrial rhythm. More broadly, they suggest that lineage-specific genome architecture can be reinforced by tissue-restricted proteins not traditionally viewed as chromatin regulators, expanding the molecular repertoire that maintains genome structure and physiological stability.

## Results

### RPL3L is a muscle-enriched ribosomal protein linked to atrial fibrillation

RPL3L is a muscle-specific ribosomal protein that has emerged as a recurrent atrial fibrillation (AF) risk locus in human genetic studies (12, 13, 14, 15, 16). To define its tissue distribution and regulatory control, we analyzed bulk and single-nucleus RNA sequencing datasets from mouse and human tissues. Across organs, *RPL3L* expression was highly enriched in striated muscles, including the heart, skeletal muscle, and tongue, and was largely absent from non-muscle tissues (Figures S1A and S1B) (17, 18). In the atrium, *RPL3L* expression was largely restricted to cardiomyocytes, with minimal expression detected in other cell populations (Figures 1A–D) (19, 20). These observations identify RPL3L as a muscle-enriched ribosomal paralog and suggest a specialized role in atrial physiology.

**Figure 1.**
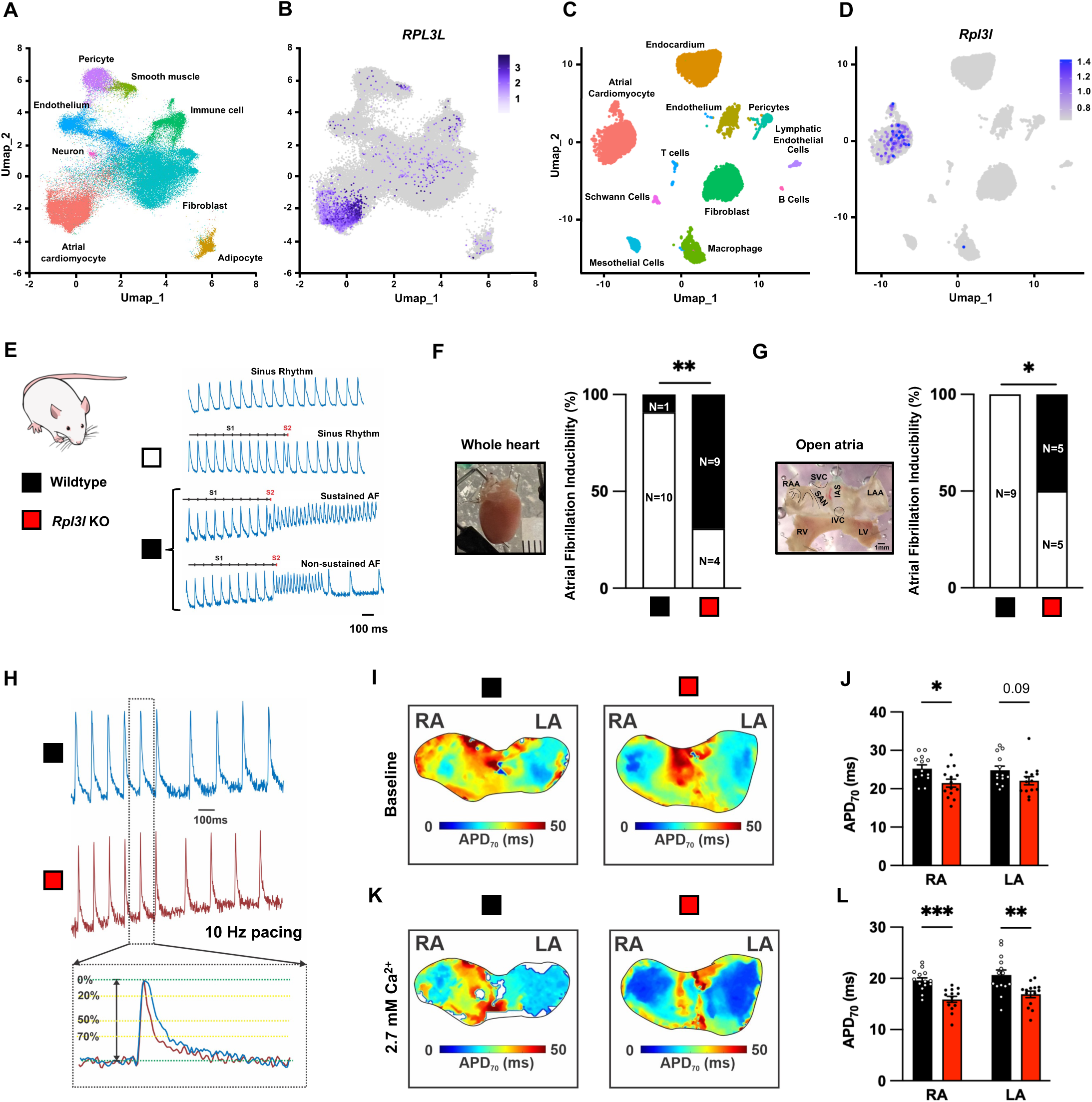
Rpl3l is muscle enriched and regulates atrial electrophysiology. (A–D) Single-nucleus RNA sequencing analysis of human and mouse hearts. UMAP visualization of major cardiac cell populations in human heart nuclei (A) and feature plot showing atrial cardiomyocyte enriched expression of *RPL3L* (B). UMAP visualization of major cardiac cell populations in mouse heart nuclei (C) and feature plot showing atrial cardiomyocyte enriched expression of *Rpl3l* (D). (E) Representative optical action potential traces of sinus rhythm, non-sustained AF, and sustained AF following S1-S2 stimulation are shown. (F and G) Quantification of AF inducibility in intact heart preparations (F) and open-atria preparations (G). Numbers within bars indicate the number of animals in each response category shown in Figure 1E. (H) Representative optical action potentials recorded during 10-Hz pacing. (I-L) Representative APD_70_ maps (I and K) and quantification (J and L) obtained from open-atria preparations under baseline conditions (I and J) and 2.7 mM CaCl_2_ (K and L) in wildtype and *Rpl3l* KO mice. Color scale indicates APD_70_ in milliseconds. Data are shown as mean ± SEM. Each dot represents an individual mouse. *\*P* < 0.05, *\*\*P* < 0.01, and *\*\*\*P* < 0.001.

Because AF is frequently associated with thyroid dysfunction (21), we next investigated whether thyroid hormone signaling regulates *Rpl3l* expression. Reanalysis of our previous datasets revealed reduced *Rpl3l* expression upon inhibition of thyroid hormone signaling (Figures S1C and S1D) (22). Inspection of publicly available chromatin profiling datasets further revealed co-occupancy of the *Rpl3l* locus by thyroid hormone receptor α (TRα) and the atrial transcription factor Tbx5 (Figures S1E and S1F), suggesting a potential atrial regulatory module (19, 22, 23, 24). Consistent with this model, *Rpl3l* expression was significantly reduced in *Myh6*-Cre; *Thra*^DN/+^ hearts during postnatal development and adulthood, as well as in pharmacological and conditional hypothyroid models induced by propylthiouracil treatment (Figures S1G–S1I). Together, these findings establish Rpl3l as a thyroid hormone-responsive gene in atrial cardiomyocytes.

To explore whether altered *RPL3L* expression may contribute to AF susceptibility, we examined AF-associated variants identified by genome-wide association studies. Analysis of GTEx datasets revealed 11 linked variants within the putative promoter region of *RPL3L* that were each associated with reduced *RPL3L* expression (Figure S1J) (25). Linkage disequilibrium analysis across the *RPL3L–NDUFB10–RPS2* locus demonstrated strong correlation among these variants, consistent with a shared haplotype block (Figure S1K). These findings suggest that reduced *RPL3L* expression may represent a common mechanism linking genetic variation at this locus to AF risk.

### *Rpl3l* deficiency increases susceptibility to atrial fibrillation

To determine whether Rpl3l is required for atrial electrical stability, we generated a *Rpl3l* knockout (KO) mouse line using CRISPR–Cas9 (Figures S2A-C). *Rpl3l* KO mutants exhibited normal cardiac function and heart size under basal conditions (Figures S2D-K), indicating that loss of *Rpl3l* does not overtly impair cardiac development or structural integrity.

Because AF is intrinsically difficult to induce in mice owing to their small atrial size, limited fibrosis, and strong electrical coupling (26), we challenged isolated hearts with elevated extracellular Ca^2+^ (2.7 mM) together with S1/S2 and burst pacing protocols (27, 28). Under these conditions, *Rpl3l* KO atria exhibited markedly increased susceptibility to AF compared with wild-type controls (Figures 1E–1G). AF was also readily induced in open-atria preparations, in which the native three-dimensional atrial architecture is disrupted, supporting an atrial-intrinsic arrhythmogenic phenotype caused by *Rpl3l* deficiency (Figure 1G).

To characterize the electrophysiological basis of this phenotype, we measured action potential duration (APD) in open-atria preparations (Figure 1H). *Rpl3l*-deficient atria exhibited a modest reduction in APD_70_ at baseline (Figures 1I and 1J). Following Ca^2+^ challenge, APD shortening became significantly more pronounced in KO atria than in controls (Figures 1K and 1L). Together, these findings indicate that Rpl3l is required to maintain atrial electrical stability and that its loss sensitizes atria to Ca^2+^-dependent electrophysiological remodeling and AF.

### Rpl3l is nuclear enriched and selectively represses the T-type calcium channel gene *Cacna1h*

Given its annotation as a ribosomal protein and prior reports of its localization to ribosomal fractions (29, 30, 31), we first examined the subcellular localization of Rpl3l in atrial tissue. Unexpectedly, immunostaining using a previously validated Rpl3l antibody (32) revealed prominent nuclear localization in atrial cardiomyocytes (Figure 2A). To independently validate this observation, we generated a C-terminal FLAG knock-in mouse line (Rpl3l-FLAG) by inserting a FLAG epitope immediately upstream of the endogenous stop codon (Figures S2L-S2N). Biochemical fractionation of atrial tissue followed by immunoblotting demonstrated enrichment of Rpl3l-FLAG in nuclear fractions relative to cytosolic fractions (Figures 2B and 2C), suggesting a previously unrecognized nuclear function for Rpl3l.

**Figure 2.**
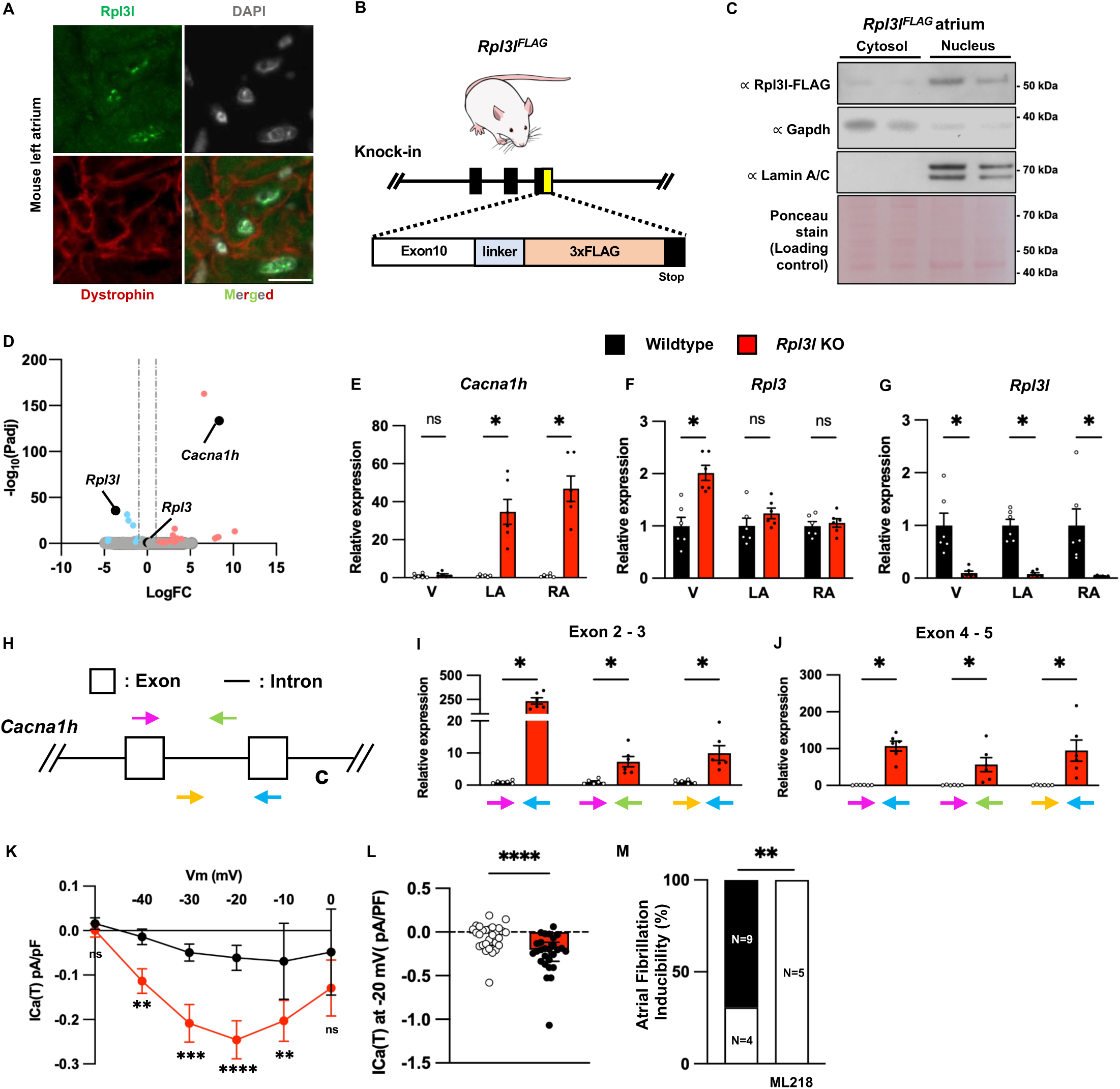
*Rpl3l* deficiency increases susceptibility to atrial fibrillation through upregulation of T-type Ca²_⁺_ channel Cacna1h. (A) Representative immunofluorescence images showing Rpl3l localization in mouse left atrium. (B) Schematic of the *Rpl3l^FLAG^* knock-in allele. A 3×FLAG tag was inserted at the C terminus of *Rpl3l*, downstream of exon 10. (C) Subcellular fractionation followed by immunoblot analysis of *Rpl3l^FLAG^* in cytosolic and nuclear fractions from atria of *Rpl3l^FLAG^* mice. Gapdh and Lamin A/C were used as cytosolic and nuclear markers, respectively. Ponceau S staining is shown as a loading control. (D) Volcano plot showing differentially expressed genes in *Rpl3l* KO left atria compared with wildtype left atria. Upregulated and downregulated genes are highlighted in orange and blue, respectively. (E–G) RT-qPCR analysis of *Cacna1h* (E), *Rpl3* (F), and *Rpl3l* (G) expression in ventricle (V), left atrium (LA), and right atrium (RA) from wildtype and *Rpl3l* KO mice. (H) Schematic of the *Cacna1h* genomic region indicating exon–intron organization and primer locations used for transcript analysis. (I and J) RT-qPCR analysis of *Cacna1h* transcripts using primer pairs spanning exon 2–3 (I) or exon 4–5 (J). (K) Whole-cell voltage-clamp recordings of T-type Ca²⁺ current density, *I*Ca(T), across the indicated membrane potentials in wildtype and *Rpl3l* KO atrial cardiomyocytes. N=28 and 27 cells from 3 wildtype and 3 KO mice, respectively. (L) Quantification of *I*Ca(T) density at −20 mV. N=28 and 27 cells from 3 wildtype and 3 KO mice, respectively. (M) Quantification of AF inducibility in ML218-treated *Rpl3l* KO hearts compared with *Rpl3l* KO hearts from Figure 1F. Numbers within bars indicate the number of animals in each category shown in Figure 1E. Data are shown as mean ± SEM. Each dot represents an individual mouse. Scale bar, 10 μm. *\*P* < 0.05, *\*\*P* < 0.01, *\*\*\*P* < 0.001, and *\*\*\*\*P* < 0.0001; ns, not significant.

To identify genes regulated by nuclear Rpl3l, we initially performed targeted qPCR analyses of ion-channel and Ca^2+^-handling genes implicated in AF. Among the candidates examined, *Cacna1h*, which encodes the T-type calcium channel Cav3.2, showed the most robust and reproducible upregulation in *Rpl3l* KO atria (Figures S3A–S3C). This observation suggested a potential link between Rpl3l and atrial Ca^2+^ homeostasis and prompted transcriptome-wide analysis.

Bulk RNA-seq confirmed selective upregulation of *Cacna1h* in KO atria, with minimal changes in other Ca^2+^-handling genes (Figure 2D). Likewise, other AF-associated ion channels and regulatory genes remained largely unchanged (Figure S3D and S3E), indicating that loss of *Rpl3l* does not broadly alter atrial electrophysiological gene programs. Follow-up qPCR analyses in an expanded cohort confirmed increased *Cacna1h* expression in KO atria but not ventricles (Figures 2E–2G). In contrast, expression of *Rpl3*, the ubiquitously expressed paralog of Rpl3l, was selectively increased in ventricular tissue, consistent with previous reports (29, 30, 31). These findings identify *Cacna1h* as a highly selective transcriptional target of Rpl3l in the atrium.

To determine whether this regulation occurs at the transcriptional level, we measured *Cacna1h* pre-mRNA using two independent primer sets. All assays revealed increased pre-mRNA abundance in KO atria (Figures 2H–2J), indicating that Rpl3l regulates *Cacna1h* transcription or cotranscriptional processing.

We next examined the functional consequences of *Cacna1h* derepression. Whole-cell voltage-clamp recordings demonstrated significantly increased T-type calcium current in isolated *Rpl3l*-deficient atrial cardiomyocytes (Figures 2K and 2L). This additional Ca^2+^ influx was associated with a significant elevation of diastolic Ca^2+^ levels in KO atrial myocytes compared with wild-type controls (Figure S3F). To determine whether and potentially how this altered Ca^2+^ homeostasis contributes to the shortened APD characteristic of *Rpl3l* deficiency, we inhibited CaMKII, a major downstream effector of Ca^2+^ signaling. Treatment with KN93 substantially prolonged APD in KO atria, restoring it toward wild-type levels (Figures S3G and S3H). These results suggest that *Cacna1h*-encoded channel-mediated Ca^2+^ entry promotes CaMKII-dependent electrophysiological remodeling in the absence of *Rpl3l*.

To determine whether the derepressed *Cacna1h*-encoded channel contributes to the pro-arrhythmic phenotype, we assessed AF inducibility in the presence of the T-type calcium channel inhibitors ML218 (33) and nickel chloride (34, 35). Both agents significantly reduced AF inducibility in *Rpl3l*-deficient atria (Figures 2M and S3I). Together, these findings demonstrate that derepression of *Cacna1h* introduces a physiologically small but functionally important T-type calcium current that elevates diastolic Ca^2+^ levels, enhances CaMKII-dependent APD shortening, and thereby increases susceptibility to AF following *Rpl3l* loss.

### Rpl3l stabilizes local chromatin insulation across the *Rpl3l*–*Cacna1h* region

To investigate how Rpl3l represses *Cacna1h* expression, we first examined whether Rpl3l directly associates with the *Cacna1h* locus. CUT&RUN analysis using Rpl3l-FLAG mice did not detect Rpl3l binding at the *Cacna1h* locus. Instead, Rpl3l was enriched at its own genomic locus, which is approximately 0.8 Mb from the *Cacna1h* locus (Figures 3A and 3B). Therefore, we hypothesized that Rpl3l regulates *Cacna1h* remotely possibly through higher-order chromatin organization rather than direct promoter occupancy.

**Figure 3.**
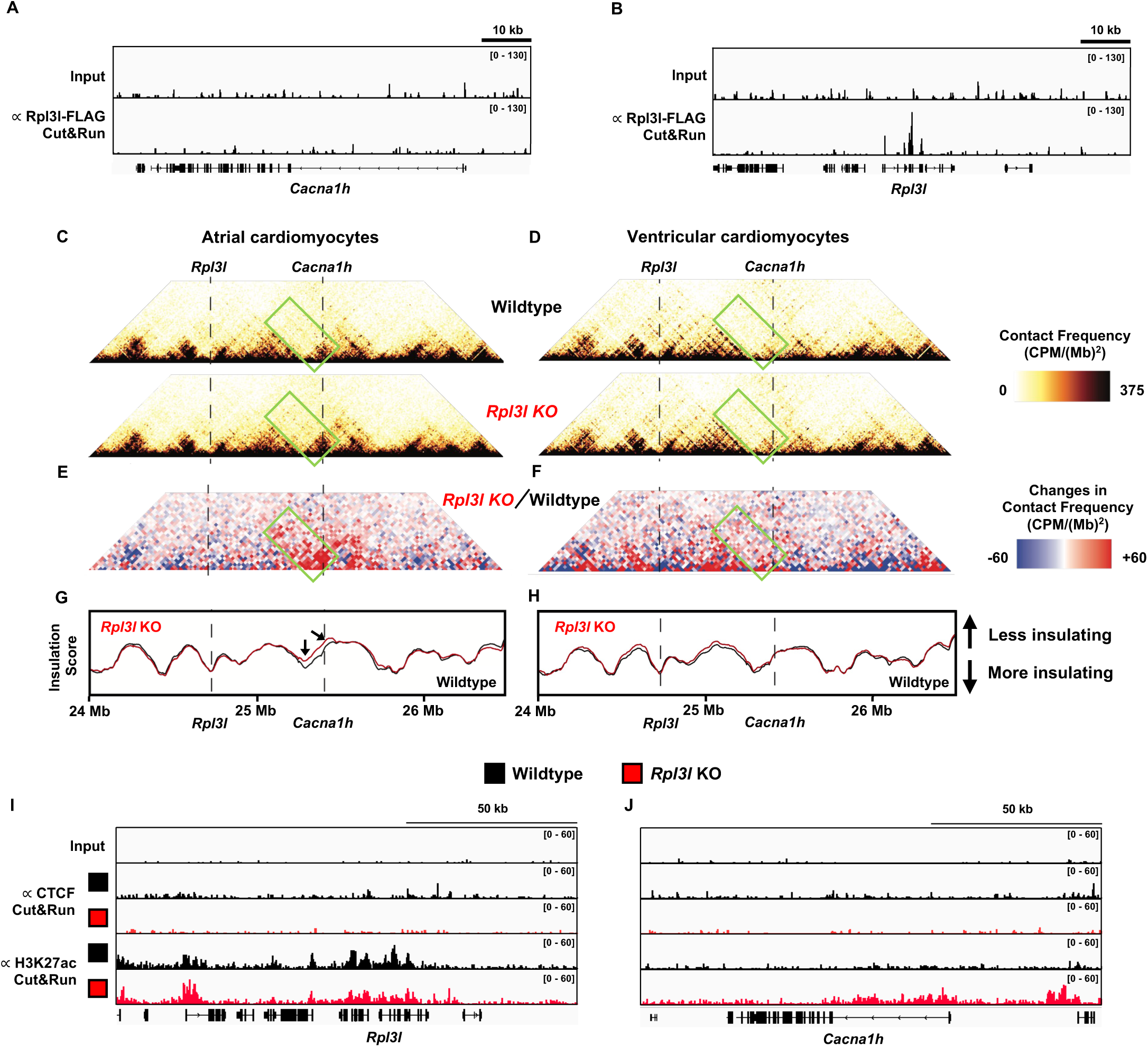
Nuclear Rpl3l regulates chromatin organization at the *Cacna1h* locus in atrial cardiomyocytes. (A and B) Genome browser tracks showing CUT&RUN signal for *Rpl3l^FLAG^* in atrial cardiomyocytes at the *Cacna1h* locus (A) and the *Rpl3l* locus (B). Input tracks are shown as control. (C and D) Hi-C contact maps from wildtype and *Rpl3l* KO cardiomyocytes across the *Rpl3l–Cacna1h* genomic region in atrial cardiomyocytes (C) and ventricular cardiomyocytes (D). Dashed lines mark the positions of the *Rpl3l* and *Cacna1h* loci. The region of increased chromatin contacts in *Rpl3l* KO is highlighted with a green box. (E and F) Differential Hi-C contact maps showing changes in chromatin contact frequency between wildtype and *Rpl3l* KO cardiomyocytes in atrial cardiomyocytes (E) and ventricular cardiomyocytes (F). Red indicates increased contact frequency in *Rpl3l* KO cardiomyocytes, and blue indicates decreased contact frequency. The region of increased chromatin contacts in *Rpl3l* KO is highlighted with a green box. (G and H) Insulation score profiles across the *Rpl3l–Cacna1h* region in atrial cardiomyocytes (G) and ventricular cardiomyocytes (H). Black and red lines indicate wildtype and *Rpl3l* KO cardiomyocytes, respectively. Reduced insulation near the *Cacna1h* locus is observed in *Rpl3l* KO atrial cardiomyocytes, with comparatively modest changes in ventricular cardiomyocytes. Arrows indicate regions with altered chromatin contacts in *Rpl3l* KO cells. (I and J) Genome browser tracks showing CUT&RUN signal for CTCF and H3K27ac at the *Rpl3l* locus (I) and *Cacna1h* locus (J) in wildtype and *Rpl3l* KO cardiomyocytes. Input tracks are shown as control.

To test this possibility, we performed Hi-C analysis in purified adult cardiomyocytes from wild-type and *Rpl3l* KO atria and ventricles. Among the four conditions examined, increased chromatin contacts spanning the *Rpl3l–Cacna1h* region were observed exclusively in *Rpl3l*-deficient atria (Figures 3C–3F), consistent with the chamber-specific upregulation of *Cacna1h* and suggesting that Rpl3l normally restricts long-range communication between these loci. Insulation score analysis further revealed a focal loss of chromatin insulation near the *Cacna1h* locus in KO atrial cardiomyocytes, whereas no comparable change was detected in ventricular cardiomyocytes (Figures 3G and 3H). These findings indicate that Rpl3l selectively stabilizes local chromatin boundary architecture in the atrial lineage.

Consistent with this model, CUT&RUN profiling revealed selective remodeling of the chromatin landscape across the *Rpl3l–Cacna1h* region in *Rpl3l*-deficient atria. In particular, the *Rpl3l* and *Cacna1h* loci exhibited altered CTCF occupancy and increased H3K27ac, consistent with weakened boundary function and transcriptional activation, respectively (Figures 3I and 3J). These changes were not evident in ventricular cardiomyocytes. Moreover, altered CTCF occupancy was not accompanied by detectable changes in DNA methylation at the locus (Figure S4A). Together, these findings support a model in which Rpl3l functions as a noncanonical chromatin regulator that stabilizes a CTCF-anchored chromatin boundary in atrial cardiomyocytes, thereby suppressing *Cacna1h* expression.

### AF-associated RPL3L variants impair nucleolar targeting and fail to repress *Cacna1h*

Previous studies have linked several coding variants in RPL3L to atrial fibrillation (13, 14, 15, 16). Here, we focused on the highest-risk missense variant, RPL3L p.Ala75Val (rs140185678, hereafter referred to as A75V), and the splice-site variant c.1167+1G>A (rs140192228), which causes exon 9 skipping (hereafter referred to as ΔS350-M389). To test whether these variants retain the ability to regulate *CACNA1H*, we expressed wild-type and mutant Rpl3ls in human iPSC-derived atrial cardiomyocytes using adenovirus. Wild-type Rpl3l reduced *CACNA1H* expression, whereas ΔS350–M389 failed to repress CACNA1H. A75V showed a weaker repressive effect than wild-type Rpl3l, suggesting that both variants impair Rpl3l-mediated *CACNA1H* regulation to varying degrees (Figures 4A and B), indicating that both variants impair Rpl3l-mediated gene expression.

**Figure 4.**
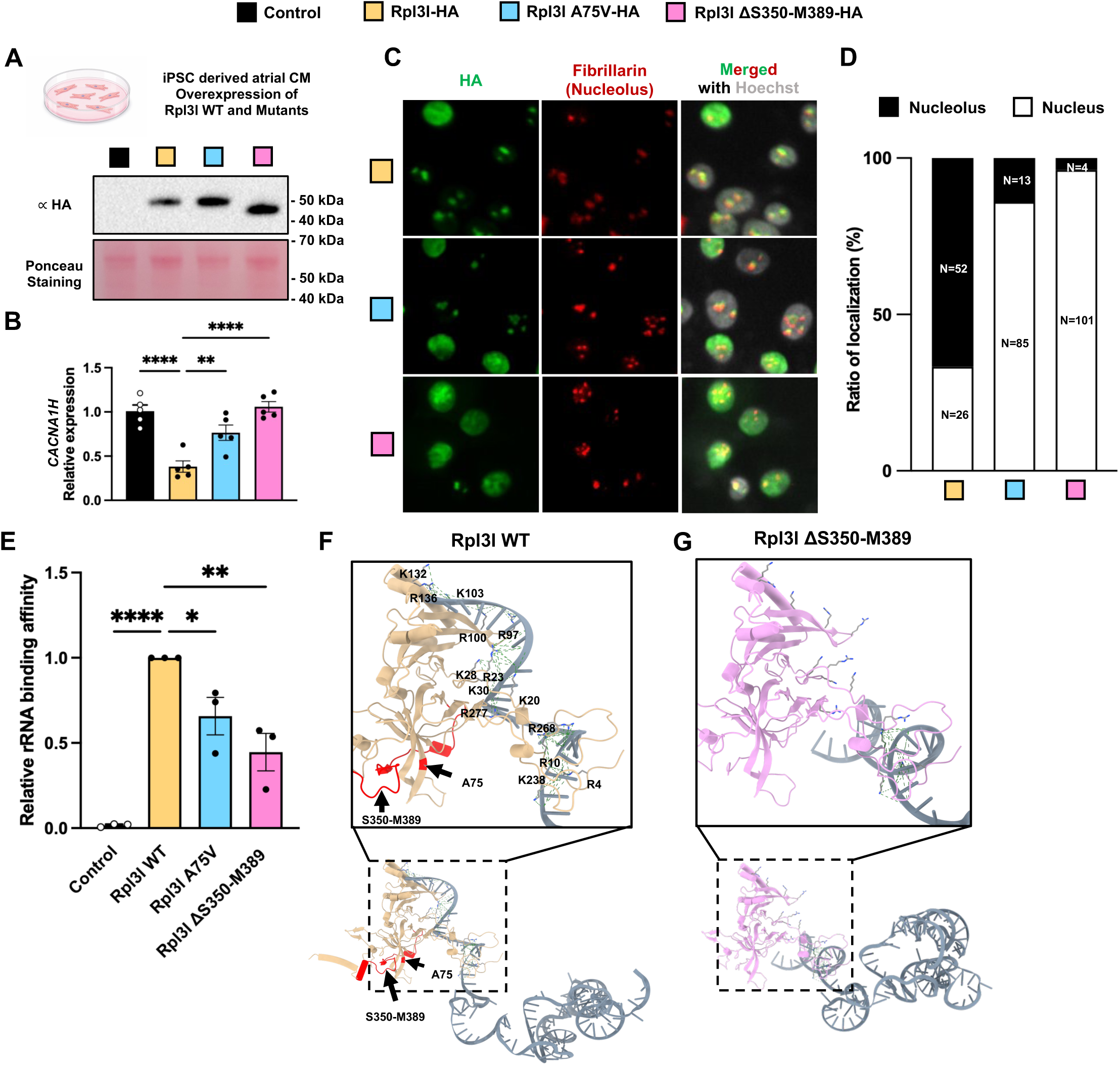
Rpl3l nucleolar localization and RNA binding contribute to regulation of *Cacna1h* expression. **(A)** Experimental schematic and immunoblot validation of Rpl3l construct expression in iPSC-derived atrial cardiomyocytes. Cells were transduced with control, Rpl3l-HA, Rpl3l A75V-HA, or Rpl3l ÄS350–M389-HA adenoviruses. Immunoblotting for HA confirms expression of Rpl3l-HA and mutant proteins. Ponceau S staining is shown as a loading control. **(B)** RT-qPCR analysis of *Cacna1h* expression in iPSC-derived atrial cardiomyocytes expressing control, Rpl3l-HA, Rpl3l A75V-HA, or Rpl3l ΔS350–M389-HA constructs. N=5 wells from one experiment. The result shown is representative of four independent experiments. **(C)** Representative immunofluorescence images showing subnuclear localization of Rpl3l-HA, Rpl3l A75V-HA, and Rpl3l ÄS350–M389-HA. HA signal is shown in green, fibrillarin-marked nucleoli are shown in red, and nuclei are counterstained with Hoechst. Merged images show overlap of HA signal with nucleolar and nuclear compartments. The result shown is representative of three independent experiments. **(D)** Quantification of Rpl3l-HA localization in nucleolar and non-nucleolar nuclear compartments. Numbers indicate the number of cells analyzed in each localization category. The result shown is representative of three independent experiments. **(E)** RNA pull-down assay measuring relative binding of Rpl3l WT, Rpl3l A75V, and Rpl3l ÄS350–M389 to ribosomal RNA. Each dot represents the RNA isolated from one animal. The result shown is representative of three independent experiments. **(F and G)** AlphaFold-predicted structural models showing RNA-proximal basic residues in Rpl3l WT **(F)** and Rpl3l ΔS350–M389 **(G)**. Lysine and arginine side chains on the predicted RNA-proximal surface are shown as sticks. Dashed green lines indicate distances of ≤10 Å between basic side-chain nitrogen atoms and RNA phosphate atoms. RNA coordinates are shown from the aligned RNA-containing structural model. The S350–M389 region and A75 residue are indicated in the WT model. Data are shown as mean ± SEM. *\*P* < 0.05, *\*\*P* < 0.01, *\*\*\*P* < 0.001, and *\*\*\*\*P* < 0.0001.

Subcellular localization analysis revealed robust nucleolar enrichment of wild-type Rpl3l, whereas both A75V and ΔS350–M389 exhibited markedly impaired nucleolar localization (Figures 4C and 4D). Because nucleolar targeting is often coupled to RNA interactions, we next examined rRNA binding. Biochemical pull-down assays demonstrated reduced rRNA association for both variants relative to wild-type Rpl3l (Figure 4E).

To investigate the structural basis of this defect, we modeled Rpl3l–RNA complexes using AlphaFold3. ΔS350–M389 produced a pronounced disruption of the predicted RNA-interacting surface, including increased separation between Rpl3l and RNA and loss of an arginine-rich helix that may contribute to RNA-backbone interactions (Figure S4B, S4C, 4F and 4G). In contrast, A75V did not substantially alter the overall predicted structure but was associated with subtle local changes near the RNA-contacting interface (Figure S4D). Together, these analyses suggest that AF-associated variants compromise Rpl3l–rRNA interactions, with ΔS350–M389 producing the more severe defect.

Collectively, these findings identify AF-associated RPL3L variants as loss-of-function alleles and indicate that nucleolar localization and rRNA binding are required for RPL3L-mediated repression of *CACNA1H*. These results establish a mechanistic link between disease-associated disruption of RPL3L subnuclear targeting and failure of ion-channel dosage control in atrial cardiomyocytes.

## Discussion

Multiple human genetic studies have identified RPL3L as one of the strongest atrial fibrillation (AF) risk loci, yet how a muscle-specific ribosomal protein contributes to arrhythmogenesis has remained unclear (13, 14, 15, 16). Unlike most ribosomal proteins, which are ubiquitously expressed, RPL3L is selectively enriched in striated muscle (36, 37), suggesting specialized functions beyond core translation. Here, we uncover an unexpected nuclear role for RPL3L in atrial cardiomyocytes and establish a mechanistic link between AF-associated genetic variation, chromatin topology, and ion-channel regulation. We show that RPL3L localizes to the nucleus and nucleolus, stabilizes a CTCF-anchored chromatin boundary across the *Rpl3l*–*Cacna1h* region, and maintains repression of the T-type calcium channel gene *Cacna1h*. Loss of this function weakens local chromatin insulation, derepresses *Cacna1h*, increases T-type calcium current, and promotes AF susceptibility. These findings identify a ribosomal protein–chromatin pathway that links three-dimensional genome organization to atrial electrical stability.

A central implication of this work concerns how differentiated cells maintain lineage-specific genome architecture. Although CTCF and cohesin establish a broadly conserved architectural framework (5, 7), chromatin insulation and long-range interactions vary substantially across tissues and developmental states (3, 38, 39). This discrepancy has suggested the existence of tissue-restricted factors that reinforce selected chromatin boundaries, yet such factors have remained largely undefined. Our data support a model in which ubiquitous architectural proteins establish potential chromatin boundaries, whereas lineage-enriched proteins stabilize selected boundaries that are particularly important for specialized cellular functions. In this context, RPL3L acts not as a global architectural factor but as a locus-selective regulator that constrains communication across the *Rpl3l*–*Cacna1h* region. More broadly, these findings suggest that cell-type-specific genome architecture emerges through cooperation between universal architectural machinery and tissue-restricted cofactors (2, 8, 9, 10, 11). RPL3L extends this concept beyond conventional DNA-binding regulators and raises the possibility that additional tissue-specific proteins contribute to chromatin insulation in differentiated cells.

The discovery that RPL3L regulates chromatin topology is particularly notable because ribosomal proteins have traditionally been viewed as structural components of the translational apparatus (40). Although several ribosomal proteins have been implicated in transcriptional regulation, RNA processing, and DNA repair (41–45), these activities have largely been characterized in cultured cells and have not been linked to higher-order genome organization in vivo. The present findings extend the functional repertoire of ribosomal proteins by demonstrating that a tissue-specific ribosomal paralog can influence chromatin architecture in a physiological and disease-relevant setting. These observations raise the possibility that other ribosomal or nucleolar proteins possess similarly specialized architectural functions in differentiated tissues.

The present study further reveals that chromatin insulation serves as a mechanism for suppressing latent developmental gene programs in adult tissues. Although *Cacna1h* is prominently expressed during cardiac development, its expression is normally maintained at low levels in the healthy adult atrium (46, 47). Our findings indicate that RPL3L-dependent boundary maintenance actively preserves this repression, thereby preventing reactivation of a developmental calcium-channel program. The physiological consequences of this topological defect are profound. Derepression of *Cacna1h* increases T-type calcium current, enhances intracellular calcium loading, and promotes AF susceptibility. Pharmacological inhibition of T-type calcium channels markedly reduced arrhythmia inducibility in *Rpl3l*-deficient atria, establishing *Cacna1h* as a key downstream effector of chromatin boundary disruption. These observations demonstrate how local changes in genome topology can be translated directly into altered cellular electrophysiology and tissue function. More broadly, they suggest that maintenance of differentiated cell identity requires continuous topological repression of developmental programs that remain encoded within the genome.

Finally, analysis of AF-associated RPL3L variants links this mechanism directly to human disease. Both A75V and ΔS350–M389 exhibited impaired nucleolar localization, reduced rRNA binding, and defective repression of *CACNA1H* in human iPSC-derived atrial cardiomyocytes. These observations suggest that AF-associated variants act as loss-of-function alleles with respect to chromatin insulation and identify nucleolar targeting as a previously unrecognized requirement for RPL3L function. In addition, these findings extend emerging evidence that nucleoli contribute to genome organization and gene regulation (48–52).

Although RPL3L variants have also been associated with cardiomyopathy in some human genetic studies (53, 54, 55, 56, 57, 58, 59), neither three previously reported *Rpl3l* knockout models nor the model examined here develop overt cardiomyopathy under basal conditions (29, 30, 31). These observations suggest that altered atrial electrical stability may represent a more primary consequence of *RPL3L* deficiency than structural myocardial dysfunction. More generally, they highlight the importance of defining tissue- and context-specific functions of ribosomal paralogs in vivo rather than inferring physiological roles solely from genetic association studies.

Together, these findings uncover a ribosomal protein–chromatin axis that couples three-dimensional genome insulation to ion-channel dosage control and atrial rhythm stability. Beyond providing a mechanistic explanation for RPL3L as an AF risk gene, this work expands current models of genome regulation by identifying a tissue-restricted ribosomal protein as a regulator of chromatin insulation. We propose that differentiated tissues employ specialized proteins not traditionally viewed as chromatin regulators to reinforce lineage-specific genome architecture and safeguard physiological function. Determining whether similar mechanisms operate in other organs, and whether additional ribosomal or nucleolar proteins contribute to chromatin topology, will be important directions for future investigation.

## Limitations of the study

This study identifies RPL3L as a noncanonical regulator of chromatin insulation and establishes *Cacna1h* derepression as a key downstream effector of *RPL3L* deficiency. However, several important questions remain unresolved. First, although *RPL3L* is expressed in both atrial and ventricular cardiomyocytes, loss of *Rpl3l* selectively disrupts chromatin insulation across the *Rpl3l*–*Cacna1h* region and derepresses *Cacna1h* in the atrium but not the ventricle. These findings suggest the existence of chamber-specific cofactors or chromatin environments that confer atrial selectivity, but the underlying mechanisms remain unknown. Second, while AF-associated RPL3L variants impair nucleolar localization, reduce rRNA binding, and fail to repress *CACNA1H* in human iPSC-derived atrial cardiomyocytes, it remains unclear whether *CACNA1H* is similarly derepressed in atrial tissue from patients carrying RPL3L risk variants or in broader subsets of human AF. Third, although our data support a role for RPL3L in stabilizing a CTCF-anchored chromatin boundary, the molecular mechanism linking nucleolar localization, rRNA interaction, and chromatin insulation remains to be defined. Addressing these questions will provide important insight into how tissue-specific proteins contribute to lineage-specific genome organization and disease susceptibility.

## Acknowledgements

We thank all members of the Huang, Lang, and Olgin laboratories, as well as the UCSF community, for helpful discussions and general support; Amanda Soe, Dr. Marco De Leon, and Hana H. Lee for preliminary experiments; Dr. Manuel Rosa-Garrido and Dr. Yuka Kitamura for advice on Hi-C; Dr. Aki Ushiki and Dr. Tatsuya Tsukui for advice on mouse generation; and Dr. Koki Tohara for advice on T-type calcium current recordings. This work is supported by the Uehara Memorial Foundation postdoc fellowship, JSPS postdoc fellowship, and PBBR postdoc independent research grant (M.N). Pershing Square, Burroughs, McKnight, and Rita Allen (L.T.). NIH (T32GM136568) and NSF (DGE-2146755) (L.C.). NIH Award (R01HL173344), AHA Career Development Award (846898), and Second Century Early Faculty Independence Award (24SCEFIA1255230) (D.L.), NIH Awards (R01HL138456 and R01HL157280), Tobacco-Related Disease Research Program Award P0558275, American Heart Association Transformative Project Award, Established Investigator Award, UCSF Program for Breakthrough Biomedical Research and BAKAR Aging Research Institute Investigator Award (G.N.H.).

## EXPERIMENTAL MODEL AND STUDY PARTICIPANT DETAILS

### Animals

All mouse and rat studies were performed in accordance with the guidelines of the University of California, San Francisco Institutional Animal Care and Use Committee (IACUC) under approved protocol AN197871. Mouse and rat strains used included C57BL/6 (Charles River Laboratories, 027); CD-1 (Charles River Laboratories, 022); Sprague Dawley (Charles River Laboratories, 001); *Myh6*-Cre (60); *TRα*^AMI/+^ (hereafter *Thra*^DN/+^) (61); a global *Rpl3l* KO line (*Rpl3l* KO, CD-1 background); and a *Rpl3l*-3×FLAG knock-in line (*Rpl3l^FLAG^*, CD-1 background). To generate the *Rpl3l* KO mice, recombinant Cas9 protein complexed with a single-guide RNA (sgRNA) targeting exon 7 of *Rpl3l* (5′-CTT GTC AGT TAC GTC GTA GC-3′) was microinjected into the pronuclei of fertilized CD-1 zygotes, which were then transferred into pseudopregnant foster females to obtain F0 founders. For the *Rpl3l^FLAG^*knock-in line, pronuclear microinjection of Cas9 protein, two sgRNAs (5′-GAG TCC TCC TTG GTG TGA AC-3′ and 5′-GTA TTA GTA GTA GCT ACC AT-3′), and a donor plasmid (backbone, pGEM-T Easy) carrying an in-frame 3×FLAG cassette was performed. For induction of hypothyroidism, CD-1 mice were fed chow containing 0.15% (w/w) propylthiouracil (PTU; Inotiv) ad libitum for 4 months; fresh chow was provided weekly.

Unless otherwise specified, adult mice were 3–6 months old, and age-matched wild-type and *Rpl3l* KO mice were used for all experiments. Both male and female mice were used unless otherwise indicated, and no sex-dependent differences were observed.

All mice were maintained in an AAALAC-accredited (Association for Assessment and Accreditation of Laboratory Animal Care) animal facility at University of California San Francisco.

### GTEx datasets and linkage-disequilibrium (LD) analysis

GTEx data were downloaded from the GTEx Portal (https://gtexportal.org/home/) and processed for downstream analysis and visualization. Linkage disequilibrium (LD) among RPL3L-associated variants was analyzed using LDmatrix through the LDlink web tool (https://ldlink.nih.gov/ldmatrix).

## METHOD DETAILS

### Echocardiography

Ventricular function was assessed by echocardiography using a Vevo 2100 ultrasound system (VisualSonics). At least three M-mode recordings were obtained from each animal, and the averaged values were used for plotting and statistical analysis.

### Optical mapping

Optical mapping was performed as previously described, with minor modifications (27, 28). Briefly, adult mouse hearts were rapidly dissected, cannulated via the aorta, and retrogradely perfused with Tyrode’s solution at a constant pressure of 80 mmHg. Hearts were loaded with the voltage-sensitive dye RH237 (Invitrogen, 0.1 mmol/L) by slow injection through a drug port over 10 min. Blebbistatin (MedchemExpress, HY-13441, 6.8 μmol/L) was subsequently administered to suppress motion artifacts. Two optical mapping preparations were used in this study: an intact heart preparation (Figure 1F) and an open-atria preparation (Figure 1G). The intact heart preparation was used for assessment of AF inducibility. For APD measurements, the atria were dissected and opened to create an open-atria preparation, allowing optical recording from the entire atrial surface. Emitted fluorescence signals were collected through >700 nm long-pass filter and acquired using a MiCAM CMOS camera (SciMedia, USA) at a sampling rate of 1 kHz and a spatial resolution of 160 μm/pixel. Atrial pacing and surface ECG were recorded using a PowerLab 26T stimulator (ADInstruments, Australia).

For AF inducibility assays, hearts were first challenged under baseline conditions and sequentially after perfusion with Tyrode’s solution containing 2.7 mM CaCl₂. Following equilibration at each condition, programmed S1–S2 stimulation and burst-pacing protocols were performed. Electrical stimulation was delivered using 1 ms pulses at twice the stimulation threshold. For the S1–S2 protocol, a train of S1 stimuli was delivered at a cycle length of 100 ms, followed by a premature S2 stimulus. The S2 coupling interval was initiated at 60 ms and progressively decreased in 2 ms steps until loss of capture or AF induction occurred. For burst pacing, hearts were paced continuously for 5 s starting at a cycle length of 60 ms, with the pacing cycle length subsequently reduced in 2 ms increments until loss of capture or AF induction. Pharmacological treatments were performed using 1 μM ML218 (MedChemExpress, HY-103309), 25 μM NiCl₂ (Sigma, 339350), or 10 μM KN93 (TargetMol, T2697), as indicated. In addition, electrical recordings were obtained during steady-state pacing at 10 Hz under all experimental conditions for action potential duration (APD) measurements and comparison.

A customized Matlab-based computer program was used to analyze the optical signals. Signals were filtered using the low-pass Butterworth algorithm at 256 Hz. Action potential duration at 70% repolarization (APD_70_) was calculated for each atrium.

### Single-cell Ca²⁺ imaging / pacing assay

Freshly isolated adult atrial cardiomyocytes from wildtype and *Rpl3l* KO mice were loaded with 1 μM Fura-2 AM (ThermoFisher, F1221) for 30 min at RT, washed, and imaged in Tyrode’s solution. Cells were electrically paced by field stimulation at 1 Hz using 5 ms pulses. Fluorescence signals were acquired using the IonOptix system at 250 frames/s. Diastolic Ca^2+^ levels were analyzed from individual rod-shaped cardiomyocytes after background correction.

### Atrial Cardiomyocyte Isolation

Single atrial cardiomyocytes were isolated for electrophysiological recording as previously described (62). Briefly, adult mice of both sexes between 12 and 16 weeks old were anesthetized with isoflurane, euthanized, and the excised heart was perfused with Tyrode’s solution containing 140 mM NaCl, 5.4 mM KCl, 1.0 mM CaCl_2_, 1.2 mM KH_2_PO_4_, 5.0 mM HEPES, 1.0 mM MgCl_2_, 5.55 mM glucose, pH7.4 with NaOH, and collagenase D+B+protease (9+15+3 mg/50 ml) via a Langendorff system perfused at 2 ml/min. After 10-15 min of digestion, the left and right atria were removed, cut into chunks, and further digested in Tyrode’s solution containing liberase (0.18 mg/ml, Roche, CA, USA) and elastase (1.0 U/ml) at 37°C for 10 min. Cells were dispersed with trituration, and a single cell suspension was collected in Kraft-Brühe solution containing: 25 mM KCl, 10 mM KH_2_PO_4_, 5 mM HEPES, 20 mM Glucose, 20 mM taurine, 100 mM K-glutamate, 10 mM K-aspartate, 2.0 mM MgSO_4_, 5.0 mM Creatine, 0.5 mM EGTA, pH 7.2 with KOH. The cell suspension was stored at 4°C, followed by at least 30 minutes at room temperature immediately before electrophysiological recording.

### Whole Cell Voltage Clamp

All cellular electrophysiology experiments were performed and analyzed by an investigator blinded to genotype. Ca^2+^ currents were recorded from 28 WT and 27 *Rpl3l* KO atrial cardiomyocytes using an Axon multiclamp-700B amplifier, Digidata 1550B and pCLAMP11.1 (Molecular Devices/Axon Instruments, Sunnyvale, CA) in whole cell voltage clamp mode. Glass microelectrodes had resistances of 2.0-2.5 MΩ. The bath solution contained (46): 140 mM TEA-Cl, 1 mM MgCl_2_, 5 mM HEPES,10 mM glucose, 3 mM CaCl_2_, 5 mM 4-aminopyridine and 0.005 mM Tetrodotoxin, pH 7.4 with CsOH. The pipette solution contained: 60 mM CsOH, 80 mM CsCl, 40 mM aspartate, 5 mM HEPES, 10 mM EGTA, 5 mM MgATP, 5 mM Na_2_-Creatine phosphate and 0.65 mM CaCl_2_ (pH 7.2 with CsOH). Calcium currents were recorded at room temperature. To elicit total and L-type Ca^2+^ currents, cells were held at -80 mV and -50 mV, respectively, and a series of test potentials was applied from -60 mV to 60 mV in 10 mV increments. Current amplitude was defined as the difference between the peak inward current and the steady-state current recorded at the end of the test pulse. Current density was calculated by normalizing the current amplitude to the cell capacitance as measured with a test pulse. T-type Ca^2+^ current tracings and amplitudes were obtained by subtracting the L-type current from the total current at each test potential.

### iPSC-derived cardiomyocyte differentiation

WTC iPSCs were differentiated into atrial cardiomyocytes using a retinoic acid-based protocol adapted from a previously established method (63). Human iPSCs were plated onto Geltrex-coated 6-well plates in Essential 8 medium (Thermo Fisher Scientific, A1517001) supplemented with 10 μM Y-27632 (MedchemExpress, HY10583) and differentiated when cultures reached 90%–100% confluence. Mesoderm induction was initiated on day 0 using RPMI+GlutaMAX (Thermo Fisher Scientific, 72400047) / B27 minus insulin (Thermo Fisher Scientific, A1895601) medium containing 8 μM CHIR99021 (Cayman Chemical, 13122), followed by Wnt inhibition with 5 μM IWP-2 (Fisher Scientific, 353310) on day 2. For atrial specification, 0.5 μM retinoic acid (Sigma, R2625) was added directly to the IWP-2-containing medium on day 3. Medium was then refreshed with RPMI+GlutaMAX/B27 (Thermo Fisher Scientific, 17504044)-based medium containing 0.5 μM retinoic acid on day 5 and maintained until day 6. Cells were maintained in RPMI+GlutaMAX/B27-based medium thereafter, replated on days 16–18, allowed to recover for 4-6 days, and purified by lactate selection for 4–6 days. Purified atrial cardiomyocytes were maintained in the recovery medium and harvested or cryopreserved on days 24–30. For transduction assays, purified atrial cardiomyocytes were plated in 96-well plates and infected with adenovirus at 1 × 10⁶–1 × 10⁷ viral genomes per well for 7 days, with medium replaced every other day.

### Bulk Easy Dip-C

Atrial and ventricular cardiomyocyte nuclei were isolated from wild-type and *Rpl3l* KO mice and processed using bulk Easy Dip-C, as described previously (64), with modifications based on the Arima-HiC kit workflow (Arima Genomics, 202507-5769). For atrial samples, 170,000 nuclei pooled from five wild-type mice and 120,000 nuclei pooled from six *Rpl3l* KO mice were used. For ventricular samples, 1,000,000 nuclei pooled from four mice were used for each genotype. Briefly, samples were crosslinked with formaldehyde to preserve chromatin contacts, and nuclei were isolated. Chromatin was digested using the Arima restriction enzyme cocktail, and digested ends were labeled with biotinylated nucleotides followed by proximity ligation under conditions favoring ligation of spatially adjacent DNA fragments. Crosslinks were reversed, and DNA was purified and mechanically sheared using a Covaris S220 sonicator with the following settings: peak power, 50.0; duty factor, 10.0; cycles per burst, 200; average power, 5.0. Samples were sheared for approximately 200 s to obtain an average fragment size of ∼400 bp. Biotinylated ligation products were captured with streptavidin beads and used for sequencing library construction, including end repair, adaptor ligation, and PCR amplification. Library quality and size distribution were assessed by shallow sequencing using a MiSeq i100. Libraries that passed quality control were sequenced on a NovaSeq X Plus platform. Hi-C reads were mapped to the mouse genome (mm10), filtered for valid ligation products, and used to generate normalized contact matrices, insulation scores, and locus-specific contact maps across the *Rpl3l*–*Cacna1h* region.

### CUT&RUN

Atrial cardiomyocyte nuclei were isolated from *Rpl3l*-*FLAG*, wild-type, and *Rpl3l* KO mice, as appropriate for each experiment. Approximately 4,000–5,000 cells were immobilized on concanavalin A–coated magnetic beads and permeabilized according to the manufacturer’s protocol (Cell Signaling Technology, 86652). Samples were incubated with primary antibodies against FLAG (Sigma, A8592), CTCF (Sigma, 07-729), H3K27ac (Cell signaling technology, 8173T), or control IgG (Cell Signaling Technology, 66362) overnight at 4°C. After washing, samples were incubated with protein A/G–MNase, and targeted chromatin cleavage was activated by the addition of calcium. Released DNA fragments were collected after incubation and purified using standard DNA purification methods. CUT&RUN libraries were prepared from purified DNA using low-input sequencing library preparation reagents compatible with Illumina sequencing and NEBNext Multiplex Oligos for Illumina (NEB, E7335S). Libraries were amplified by PCR, assessed for quality and size distribution, and sequenced using paired-end reads. Sequencing reads were mapped to the mouse genome (mm10), and low-quality reads, PCR duplicates, and mitochondrial reads were removed. Genome-wide enrichment profiles were generated and normalized across samples. Peaks were called for each antibody, and signal tracks were visualized across the *Rpl3l*–*Cacna1h* region.

### RNA-seq

Total RNA was extracted from mouse left atrial tissues using TRIzol reagent (Invitrogen, 15596-018) according to standard RNA isolation procedures. RNA quality and concentration were assessed before library preparation. RNA-seq library preparation, sequencing, and the downstream analyses were performed by Novogene. Sequencing reads were aligned to the mouse reference genome (mm10), and gene-level counts were generated using annotated gene models. Low-quality reads and reads mapping to multiple genomic locations were excluded from downstream analyses. Differential gene expression analysis was performed between wild-type and *Rpl3l* KO samples.

### Bisulfite sequencing

Genomic DNA was extracted from isolated mouse atrial cardiomyocytes using the Quick-DNA Miniprep Kit (Zymo, D4068). Bisulfite conversion was performed using the EZ DNA Methylation-Gold Kit (Zymo, D5005) according to the manufacturer’s instructions. Primer sets were designed to amplify four loci surrounding the CTCF-binding site. PCR amplification was performed using PCRBIO HS-Taq polymerase (PCR Biosystems, PB10.22-02), and the resulting amplicons were cloned into a TOPO vector (Thermo Fisher Scientific, 450030). Individual clones were then sequenced by nanopore sequencing (Quintara).

### rRNA pull down assay

GST-fusion proteins comprising wild-type Rpl3l, Rpl3l A75V, and Rpl3l ΔS350–M389 were expressed in *Escherichia coli* BL21 and purified using glutathione agarose beads (Sigma, G4510). Total RNA was isolated from cardiac tissues, and 2–3 μg of total RNA was pre-cleared with GST-bound glutathione agarose beads for 10 min before pull-down. Pre-cleared RNA was incubated with equal amounts (∼1 μg) of GST, GST-Rpl3l, GST-Rpl3l A75V, or GST-Rpl3l ΔS350–M389 in the presence of RNase inhibitor (Promega, N2611) for 1 h at 4°C. Beads were washed sequentially with low-salt and high-salt wash buffers containing 150 mM and 250 mM NaCl, respectively, and bound RNA was eluted with TRIzol reagent. Recovered RNA was purified and analyzed by qRT-PCR to assess rRNA association.

### Quantitative PCR (qPCR)

Tissues were homogenized in TRIzol reagent, and total RNA was extracted according to the manufacturer’s protocol. cDNA was synthesized using the iScript cDNA Synthesis Kit (Bio-Rad), and qPCR was performed using SYBR Select Master Mix (Thermo Fisher Scientific, 4472908). Primer sequences are listed in the supplemental table.

### Western Blotting

Nuclear and cytoplasmic fractions were separated using the Nuclear and Cytoplasmic Extraction Reagents (Thermo Fisher Scientific, 78833) according to the manufacturer’s instructions. Protein concentrations were measured using a BCA Protein Assay Kit (Thermo Fisher Scientific, 23227). For protein separation, 15-30 μg of protein lysate was loaded onto a NuPAGE 4-12% Bis-Tris Gel (Invitrogen, NP0335 and NP0336), followed by transfer to a PVDF membrane (Bio-Rad, 1620177). Protein loading was assessed by Ponceau S staining (VWR, 97063-650). The membrane was then blocked with 5% milk in TBS containing 0.1% Triton X-100 and incubated with the indicated antibodies. Blots were visualized using ECL Western Blotting Substrate (Genesee, 20-300). To reprobe the membrane with different antibodies, the membrane was stripped three times for 10 min each with 10 mM Tris, 150 mM NaCl, pH 2.3, then blocked again and incubated with antibodies. The following primary antibodies were used: rabbit anti-HA-HRP antibody (Roche, 12013819001), mouse anti-Gapdh-HRP antibody (Abclonal, AC035), rabbit anti-LaminA/C antibody (Abclonal, A0249); mouse FLAG antibody (Sigma, F3165), mouse HRP IgG (GENA931), rabbit HRP IgG (GENA934).

### Histology and immunofluorescence staining

At the indicated time point, animals were anesthetized by intraperitoneal injection of 10% ethyl carbamate in 1× PBS. Hearts were rapidly excised, rinsed to remove blood, briefly incubated in 30% sucrose in 1× PBS, embedded in O.C.T. Compound (Tissue-Tek, cat#4583), and snap-frozen on dry ice. Hearts were cryosectioned at 5 μm onto charged slides. Sections were fixed in 4% paraformaldehyde (PFA) in 1× PBS, washed three times with 0.1% Triton X-100 in PBS (PBST), and blocked in 5% normal donkey serum in PBST. Primary antibodies were then applied, followed by three washes in PBST and incubation with Alexa Fluor-conjugated secondary antibodies (1:500). After three additional washes in PBST, nuclei were counterstained with DAPI. The following primary antibodies were used: Rpl3l (in-house polyclonal rabbit antibody generated by John McCarthy), rabbit anti-HA antibody (Abclonal, AE105), Alexa Fluor 488 donkey anti-rabbit IgG (Thermo, A-21206), Alexa Fluor 555 donkey anti-rabbit IgG (Thermo, A-31572).

### Viral constructs and transduction

Rpl3l coding sequences corresponding to wild-type Rpl3l, Rpl3l A75V, and Rpl3l ΔS350–M389 were cloned into adenoviral expression vectors (pAdTrack-CMV). Adenovirus was produced using standard packaging methods and used (65) to transduce purified human iPSC-derived atrial cardiomyocytes. For transduction assays, cells were plated in 96-well plates and infected with adenovirus at 1 × 10⁶–1 × 10⁷ viral genomes per well for 7 days, with medium replaced every other day. Cells were then harvested for downstream gene-expression analysis and immunofluorescence staining.

### AlphaFold3 structural modeling

Structural models of wild-type Rpl3l and Rpl3l with AF-associated variants in complex with RNA were generated using AlphaFold3. Protein sequences corresponding to wild-type Rpl3l (ENSMUST00000170239.8), Rpl3l A75V, and Rpl3l ΔS350–M389 were submitted together with a partial 28S rRNA sequence used for modeling (GUA ACC CGU UGA ACC CCA UUC GUG AUG GGG AUC GGG GAU UGC AAU UAU UCC CCA UGA ACG AGG AAU UCC CAG UAA GUG CGG GUC AUA AGC UUG CGU UGA UUA AGU CCC UGC CCU UUG UAC ACA CCG CCC GUC GCU ACU ACC GAU UGG AUG G) used for modeling. Multiple predictions were generated for each protein–RNA complex, and representative top-ranked models were used for visualization and comparison. Predicted protein–RNA interfaces were inspected in UCSF ChimeraX, and local changes in RNA-facing regions were compared among wild-type and mutant Rpl3l models.

### Statistical analysis and quantification

Statistical analyses were performed using GraphPad Prism. Data are presented as mean ± SEM unless otherwise indicated. For comparisons between two groups, the Mann–Whitney U test was used unless otherwise specified. For comparisons among multiple groups, one-way or two-way ANOVA followed by appropriate post hoc multiple-comparison tests was used. Categorical data, including AF inducibility, were analyzed using the fisher’s exact test, as appropriate. The number of biological replicates, cells, or independent experiments is indicated in the figures or figure legends. A P value of <0.05 was considered statistically significant.

## Data and code availability

RNA-seq, CUT&RUN, and Hi-C sequencing datasets generated in this study will be deposited in the Gene Expression Omnibus. Publicly available datasets used in this study are listed in the key resources table. No custom software was generated for this study. Additional information required to reanalyze the data reported in this paper is available from the lead contact upon reasonable request.

## Author contributions

M.N., D.L., and G.N.H. designed the research. M.N., X.C., S.Y., R.H., A.B., N.L., K.K., K.H., D. L., and G.N.H. performed experiments. M.N., X.C., S.Y., L.X.C., R.H., A.B., K.K., D. L., and G.N.H. analyzed data. J.P., A.M., Y.C., V.V., L.T., J.E.O., D.L., and G.N.H. provided supervision, reagents, and conceptual input. M.N., S.Y., D.L., and G.N.H. wrote the manuscript with input from all authors.

---

**Table 1.**
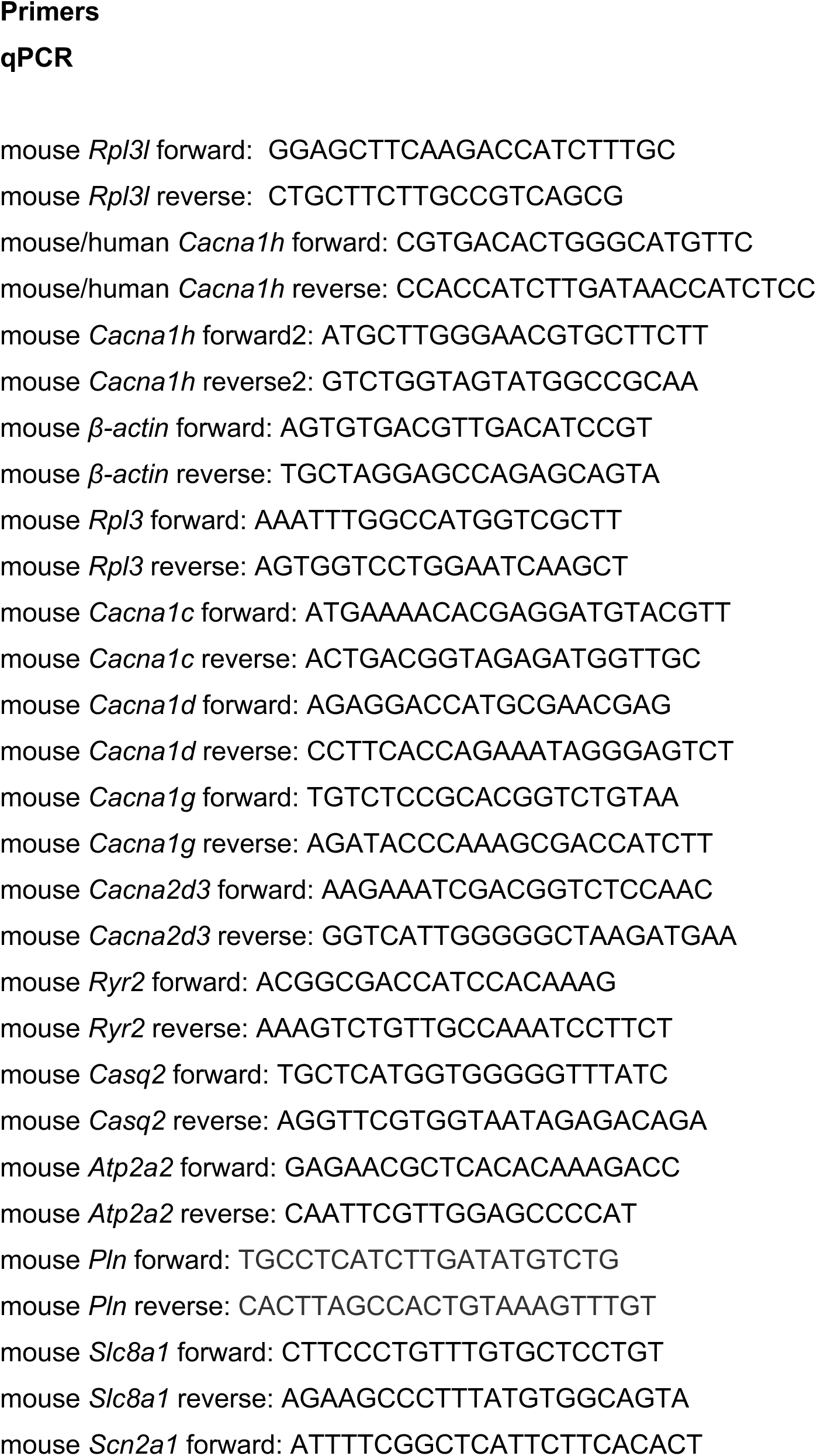
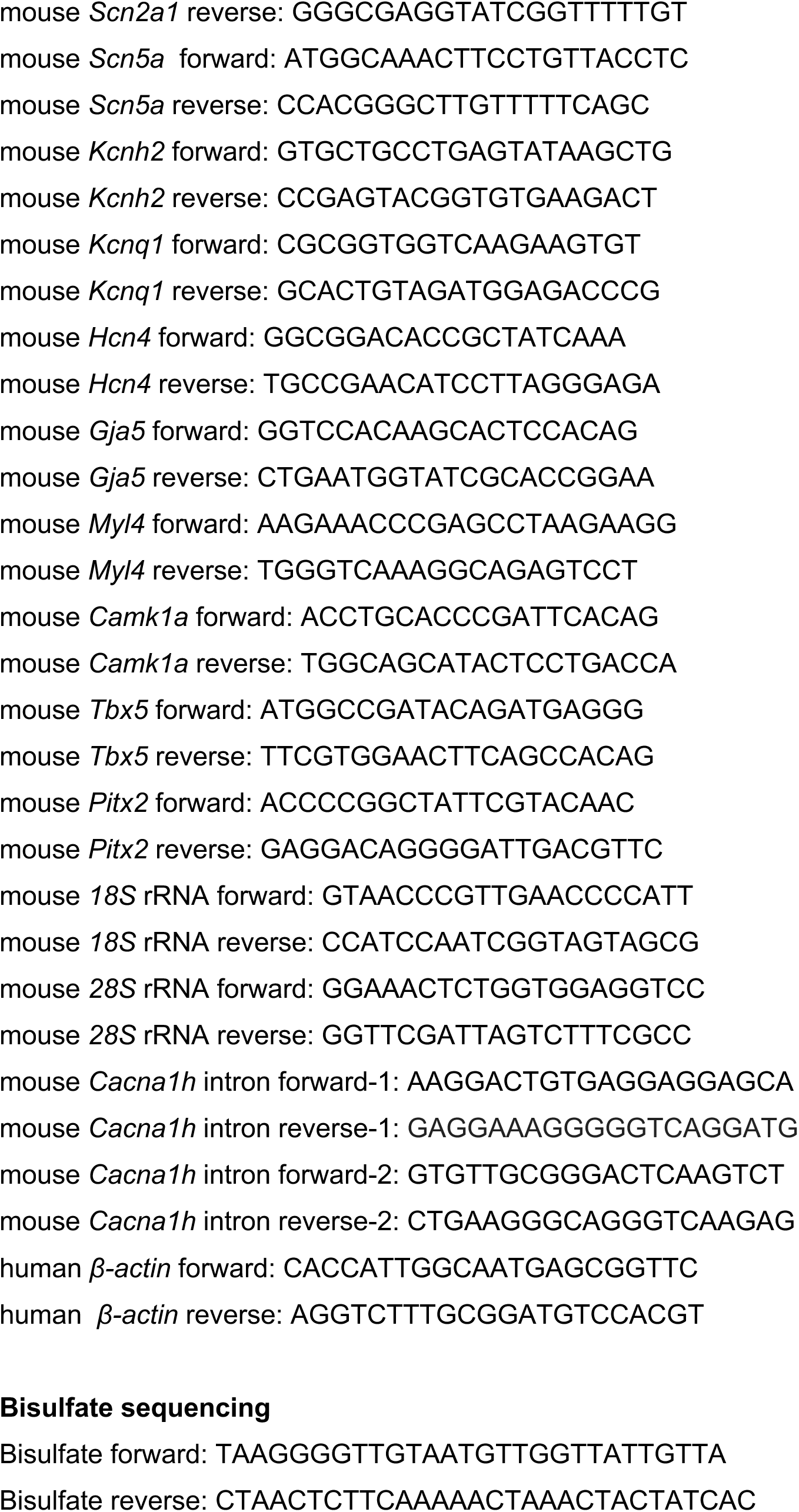
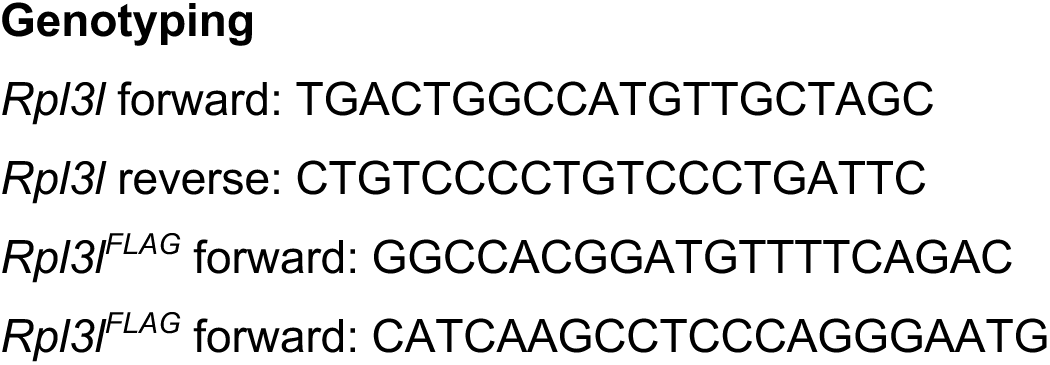

**Supplementary Figure 1.**
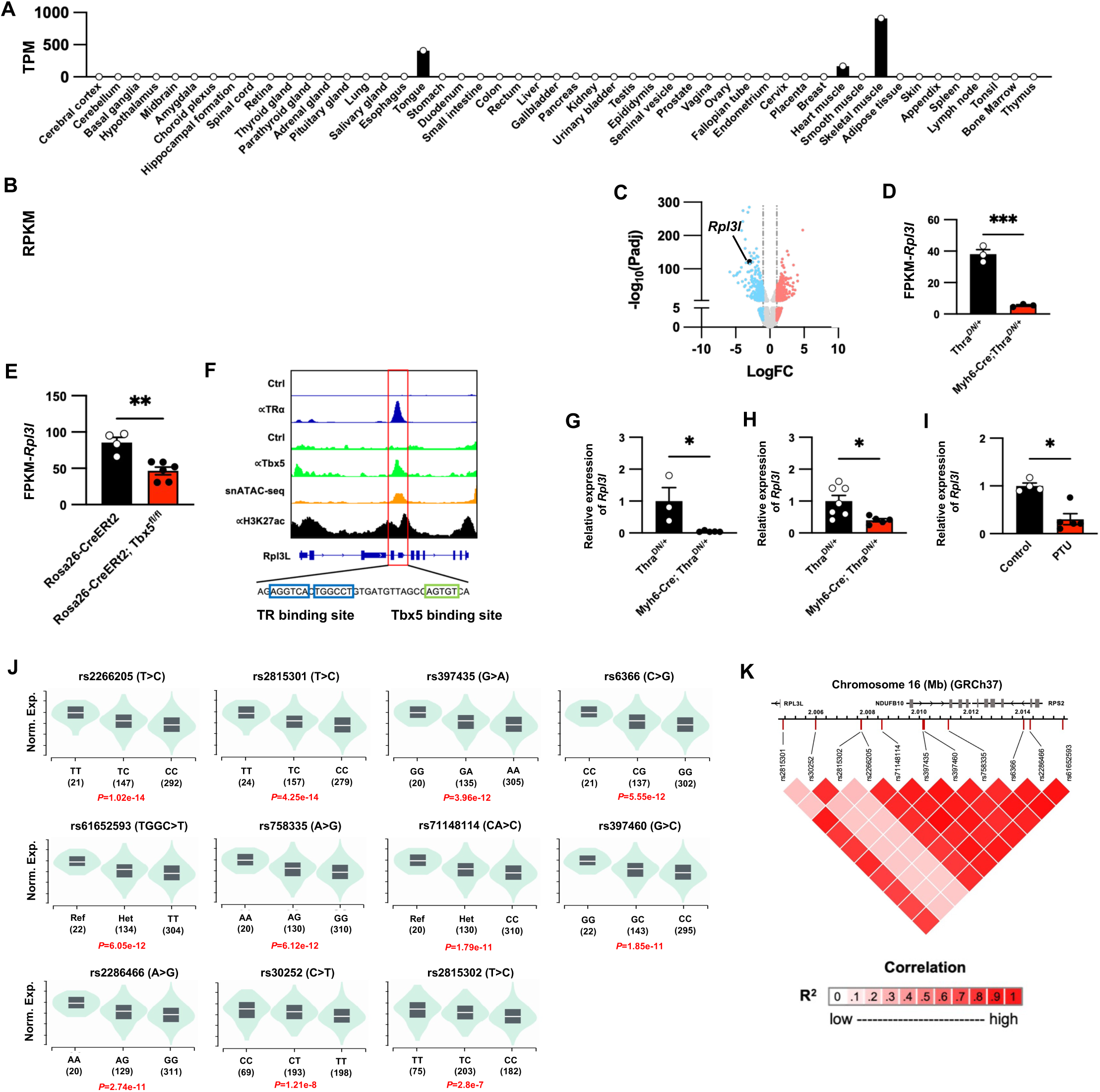
Thyroid hormone signaling and atrial fibrillation–associated genetic variation converge on *Rpl3l* regulation. (A) Tissue distribution of *RPL3L* expression across human organs from RNA-seq datasets, shown as transcripts per million (TPM). (B) Tissue distribution of *Rpl3l* expression across mouse organs from RNA-seq datasets, shown as reads per kilobase per million mapped reads (RPKM). (C) Volcano plot showing differential gene expression in *Myh6*-Cre; *Thra*^DN/+^ hearts. (D) *Rpl3l* expression in ventricles from *Thra*^DN/+^ and *Myh6*-Cre; *Thra*^DN/+^ mice. (E) *Rpl3l* expression in ventricles from Rosa26^CreERT2^ control and Rosa26^CreERT2^; *Tbx5*^fl/fl^ mice. (F) Genome browser tracks showing TRá, Tbx5, H3K27ac ChIP-seq, and snATAC-seq signal at the *Rpl3l* locus. The predicted TR and Tbx5 binding sites are indicated below the tracks. (G-H) RT-qPCR analysis of *Rpl3l* expression in P14 (G) and adult (H) left atria from *Thra*^DN/+^ and *Myh6*-Cre; *Thra*^DN/+^ mice. (I) RT-qPCR analysis of *Rpl3l* expression in adult left atria from control and PTU treated mice. (J) Distribution of *RPL3L* expression in human atrial appendage tissue from GTEx, stratified by genotype for atrial fibrillation–associated variants. Normalized expression is shown for each genotype group; nominal *P* values are indicated. (K) Linkage disequilibrium structure among variants associated with reduced *RPL3L* expression in human atrial appendage tissue. Pairwise correlation is shown as *R^2^*. Data are shown as mean ± SEM. Each dot represents an individual animal. *\*P* < 0.05, *\*\*P* < 0.01, and *\*\*\*P* < 0.001

**Supplementary Figure 2.**
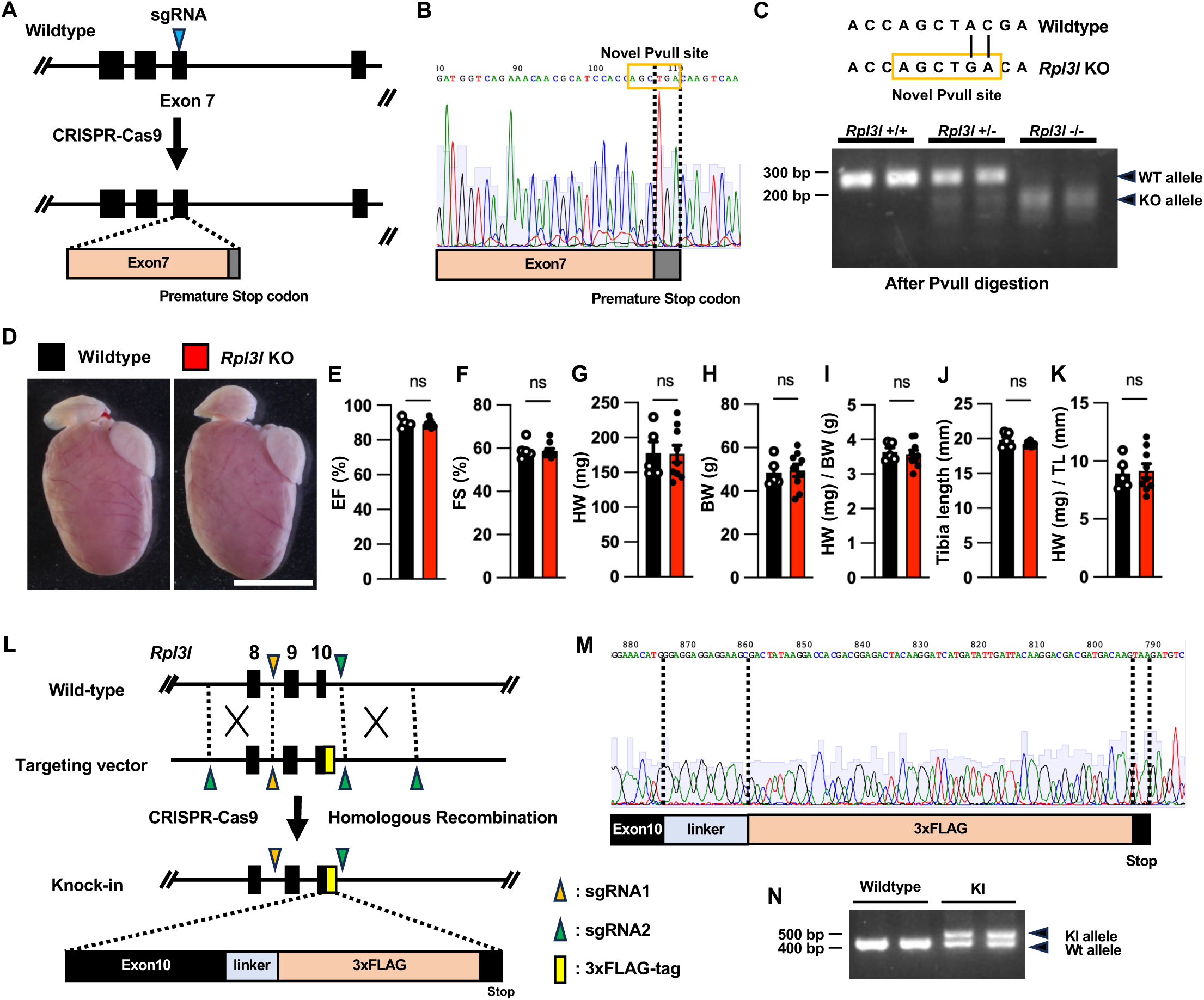
Generation and phenotypic validation of *Rpl3l* KO and *Rpl3l^FLAG^* mice. (A) Schematic of the CRISPR-Cas9 genome-editing strategy used to generate *Rpl3l* KO mice. The sgRNA was designed to target exon 7 of the *Rpl3l* locus, and homology-directed repair introduced a premature stop codon within exon 7. (B) Sanger sequencing chromatogram confirming introduction of the engineered mutant allele. The mutation introduced a premature stop codon together with a novel PvuII restriction site, CAG^CTG, within exon 7. (C) PCR-based genotyping strategy using PvuII restriction digestion. Wild-type mice show the undigested allele, heterozygous mice show both wild-type and digested mutant alleles, and homozygous *Rpl3l* KO mice show only the digested mutant allele, confirming germline transmission of the edited allele. (D) Representative gross heart images from wildtype and *Rpl3l* KO mice. (E–K) Quantification of baseline cardiac anatomical and functional parameters in wildtype and *Rpl3l* KO mice, including ejection fraction (EF) (E), fractional shortening (FS) (F), heart weight (HW) (G), body weight (BW) (H), heart weight normalized to body weight (HW/BW) (I), tibia length (J), and heart weight normalized to tibia length (HW/TL) (K). (L) Schematic of the CRISPR-Cas9 mediated knock-in strategy used to generate C-terminally tagged *Rpl3l^FLAG^* mouse. sgRNAs were designed to target the region surrounding exon 10 of the *Rpl3l* locus, and homologous recombination introduced a linker sequence followed by a 3×FLAG tag immediately downstream of exon 10. (M) Sanger sequencing chromatogram confirming correct insertion of the linker-3×FLAG cassette at the C terminus of *Rpl3l*. The exon 10, linker, 3×FLAG, and stop-codon regions are indicated. (N) PCR-based genotyping of wildtype and *Rpl3l^FLAG^* mice. Wildtype mice show the unmodified allele (400 bp), whereas *Rpl3l^FLAG^* mice show both the wildtype and knock-in alleles (500 bp), confirming successful germline transmission of the targeted allele. Data are shown as mean ± SEM. Each dot represents an individual animal. Scale bar, 1 cm. ns, not significant

**Supplementary Figure 3.**
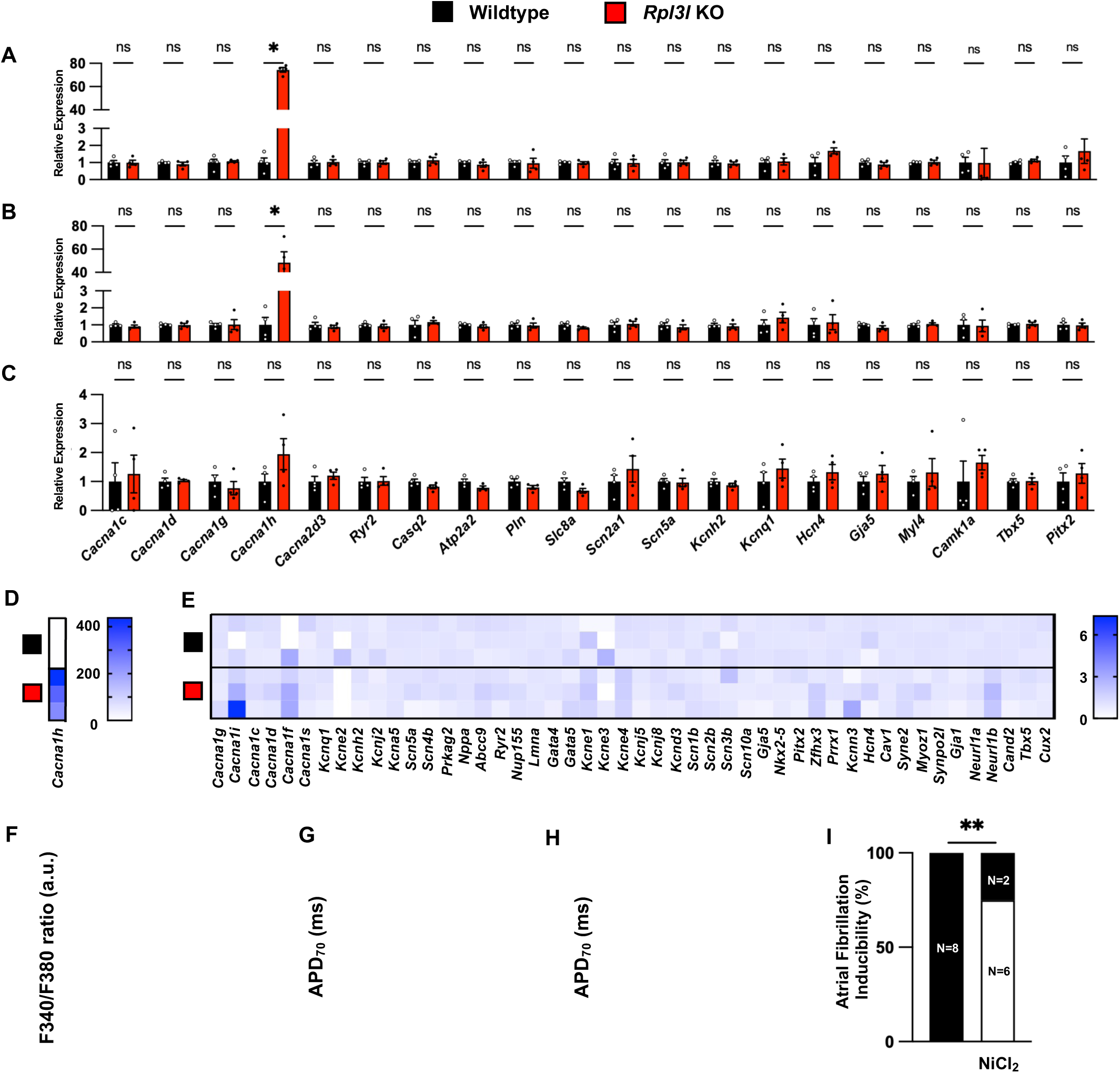
*Rpl3l* deficiency selectively derepresses *Cacna1h* and disrupts atrial Ca²_⁺_ homeostasis. (A – C) RT-qPCR analysis of selected cardiac ion-channel and calcium-handling genes in mouse wildtype and *Rpl3l* KO right atria (A), left atria (B), and ventricle (C). (D and E) Summary heatmaps showing relative expression of *Cacna1h* (D) and additional ion-channel and calcium signaling pathway genes (E) in wildtype and *Rpl3l* KO atrial samples. Color scales indicate relative expression. (F) Quantification of the F340/F380 ratio metric Ca^2^⁺ signal in wildtype and *Rpl3l* KO atria. Each dot represents an individual atrial cardiomyocyte. (G and H) Quantification of atrial action potential duration at 70% repolarization (APD_70_) following KN-93 treatment in right atria (G) and left atria (H) from wildtype and *Rpl3l* KO mice. (I) Quantification of AF inducibility with NiCl_2_. Numbers within bars indicate the number of animals in each category shown in Figure 1E. Data are shown as mean ± SEM. Each dot represents an individual animal. ns, not significant; *\*P* < 0.05, and *\*\*P* < 0.01.

**Supplementary Figure 4.**
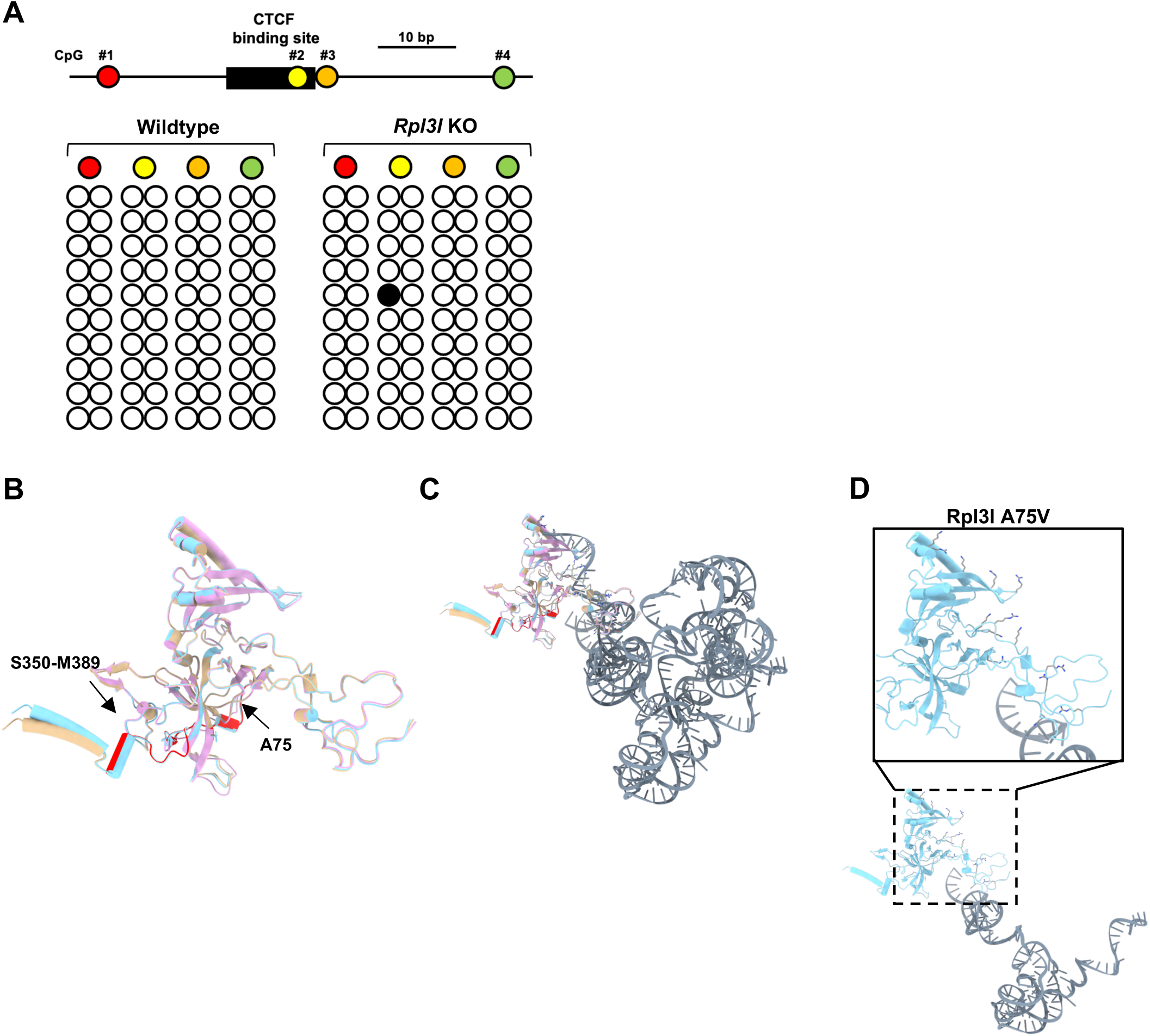
Molecular characterization of Rpl3l mutants and predicted RNA-associated structural features. (A) Bisulfite sequencing analysis of CpG methylation across the predicted CTCF-binding region. The schematic indicates the positions of CpG sites analyzed relative to the CTCF-binding site. Open circles represent unmethylated CpGs, and filled circles represent methylated CpGs in individual sequenced clones from wild-type and *Rpl3l* KO samples. (B) Structural alignment of predicted Rpl3l WT, Rpl3l A75V, and Rpl3l ΔS350–M389 models. WT, A75V, and ΔS350–M389 models are shown in tan, cyan, and magenta, respectively. The A75 residue and the S350–M389 region are highlighted in red. (C) Overview of the predicted Rpl3l structural model aligned to an RNA-containing ribosomal structural context. Rpl3l is shown in the aligned protein model colors, and RNA is shown in gray, illustrating the predicted spatial relationship between Rpl3l and nearby RNA. (D) AlphaFold predicted structural models showing RNA-proximal basic residues in Rpl3l A75V mutant.

## Notes

### Competing Interest Statement

The authors have declared no competing interest.

## References

1. Bonev, B., and Cavalli, G. (2016). Organization and function of the 3D genome. Nat. Rev. Genet. 17, 661–678. 10.1038/nrg.2016.112

2. Stadhouders, R., Filion, G.J., and Graf, T. (2019). Transcription factors and 3D genome conformation in cell-fate decisions. Nature 569, 345–354. 10.1038/s41586-019-1182-7

3. Dixon, J.R., Selvaraj, S., Yue, F., Kim, A., Li, Y., Shen, Y., Hu, M., Liu, J.S., and Ren, B. (2012). Topological domains in mammalian genomes identified by analysis of chromatin interactions. Nature 485, 376–380. 10.1038/nature11082

4. Ji, X., Dadon, D.B., Powell, B.E., Fan, Z.P., Borges-Rivera, D., Shachar, S., Weintraub, A.S., Hnisz, D., Pegoraro, G., Lee, T.I., et al. (2016). 3D chromosome regulatory landscape of human pluripotent cells. Cell Stem Cell 18, 262–275. 10.1016/j.stem.2015.11.007

5. Rao, S.S.P., Huntley, M.H., Durand, N.C., Stamenova, E.K., Bochkov, I.D., Robinson, J.T., Sanborn, A.L., Machol, I., Omer, A.D., Lander, E.S., et al. (2014). A 3D map of the human genome at kilobase resolution reveals principles of chromatin looping. Cell 159, 1665–1680. 10.1016/j.cell.2014.11.021

6. Lupiáñez, D.G., Kraft, K., Heinrich, V., Krawitz, P., Brancati, F., Klopocki, E., Horn, D., Kayserili, H., Opitz, J.M., Laxova, R., et al. (2015). Disruptions of topological chromatin domains cause pathogenic rewiring of gene-enhancer interactions. Nature 518, 363–367. 10.1038/nature14222

7. Grubert, F., Srivas, R., Spacek, D.V., Kasowski, M., Ruiz-Velasco, M., Sinnott-Armstrong, N., Greenside, P., Narasimha, A., Liu, Q., Geller, B., et al. (2020). Landscape of cohesin-mediated chromatin loops in the human genome. Nature 583, 737–743. 10.1038/s41586-020-2151-x

8. Wang, R., Chen, F., Chen, Q., Wan, X., Shi, M., Chen, A.K., Ma, Z., Li, G., Wang, M., Ying, Y., et al. (2022). MyoD is a 3D genome structure organizer for muscle cell identity. Nat. Commun. 13, 205. 10.1038/s41467-021-27865-6

9. Johanson, T.M., Lun, A.T.L., Coughlan, H.D., Tan, T., Smyth, G.K., Nutt, S.L., and Allan, R.S. (2018). Transcription-factor-mediated supervision of global genome architecture maintains B cell identity. Nat. Immunol. 19, 1257–1264. 10.1038/s41590-018-0234-8

10. Wahl, N., Espeso-Gil, S., Chietera, P., Nagel, A., Laighneach, A., Morris, D.W., Rajarajan, P., Akbarian, S., Dechant, G., and Apostolova, G. (2024). SATB2 organizes the 3D genome architecture of cognition in cortical neurons. Mol. Cell 84, 621–639.e9. 10.1016/j.molcel.2023.12.024

11. Grant, Z.L., Kuang, S., Zhang, S., Horrillo, A.J., Rao, K.S., Kameswaran, V., Joubran, C., Lau, P.K., Dong, K., Yang, B., et al. (2025). Dose-dependent sensitivity of human 3D chromatin to a heart disease-linked transcription factor. bioRxiv. 10.1101/2025.01.09.632202

12. Chaillou, T., Zhang, X., and McCarthy, J.J. (2016). Expression of muscle-specific ribosomal protein L3-like impairs myotube growth. J. Cell. Physiol. 231, 1894–1902. 10.1002/jcp.25294

13. Thorolfsdottir, R.B., Sveinbjornsson, G., Sulem, P., Nielsen, J.B., Jonsson, S., Halldorsson, G.H., Melsted, P., Ivarsdottir, E.V., Davidsson, O.B., Kristjansson, R.P., et al. (2018). Coding variants in RPL3L and MYZAP increase risk of atrial fibrillation. Commun. Biol. 1, 68. 10.1038/s42003-018-0068-9

14. Vad, O.B., Monfort, L.M., Paludan-Müller, C., Kahnert, K., Diederichsen, S.Z., Andreasen, L., Lotta, L.A., Nielsen, J.B., Lundby, A., Svendsen, J.H., et al. (2024). Rare and common genetic variation underlying atrial fibrillation risk. JAMA Cardiol. 9, 732–740.

15. Choi, S.H., Jurgens, S.J., Xiao, L., Hill, M.C., Haggerty, C.M., Sveinbjörnsson, G., Morrill, V.N., Marston, N.A., Weng, L.-C., Pirruccello, J.P., et al. (2025). Sequencing in over 50,000 cases identifies coding and structural variation underlying atrial fibrillation risk. Nat. Genet. 57, 548–562. 10.1038/s41588-025-02074-9

16. Roselli, C., Surakka, I., Olesen, M.S., Sveinbjornsson, G., Marston, N.A., Choi, S.H., Holm, H., Chaffin, M., Gudbjartsson, D., Hill, M.C., et al. (2025). Meta-analysis of genome-wide associations and polygenic risk prediction for atrial fibrillation in more than 180,000 cases. Nat. Genet. 57, 539–547. 10.1038/s41588-024-02072-3

17. Li, B., Qing, T., Zhu, J., Wen, Z., Yu, Y., Fukumura, R., Zheng, Y., Gondo, Y., and Shi, L. (2017). A comprehensive mouse transcriptomic BodyMap across 17 tissues by RNA-seq. Sci. Rep. 7, 4200. 10.1038/s41598-017-04520-z

18. The GTEx Consortium (2015). The Genotype-Tissue Expression (GTEx) pilot analysis: multitissue gene regulation in humans. Science 348, 648–660. 10.1126/science.1260419

19. Steimle, J.D., Canozo, F.J.G., Park, M., Kadow, Z.A., Samee, M.A.H., and Martin, J.F. (2022). Decoding the PITX2-controlled genetic network in atrial fibrillation. JCI Insight 7, e158895. 10.1172/jci.insight.158895

20. Tucker, N.R., Chaffin, M., Fleming, S.J., Hall, A.W., Parsons, V.A., Bedi, K.C., Jr., Akkad, A.D., Herndon, C.N., Arduini, A., Papangeli, I., et al. (2020). Transcriptional and cellular diversity of the human heart. Circulation 142, 466–482. 10.1161/CIRCULATIONAHA.119.045401

21. Singh, H., Shahid, M.Z., Harrison, S.L., Lane, D.A., Lip, G.Y.H., and Logantha, S.J.R.J. (2024). Subclinical thyroid dysfunction and the risk of incident atrial fibrillation: A systematic review and meta-analysis. PLoS One 19, e0296413. 10.1371/journal.pone.0296413

22. Hirose, K., Payumo, A.Y., Cutie, S., Hoang, A., Zhang, H., Guyot, R., Lunn, D., Bigley, R.B., Yu, H., Wang, J., et al. (2019). Evidence for hormonal control of heart regenerative capacity during endothermy acquisition. Science 364, 184–188. 10.1126/science.aar2038

23. Rathjens, F.S., Garbern, J.C., Wieland, T., Keegan, B.R., Novoyatleva, T., and Puceat, M. (2021). Preclinical evidence for the therapeutic value of TBX5 normalization in arrhythmia control. Cardiovasc. Res. 117, 1908–1922. 10.1093/cvr/cvaa211

24. Nord, A.S., Blow, M.J., Attanasio, C., Akiyama, J.A., Holt, A., Hosseini, R., Phouanenavong, S., Plajzer-Frick, I., Shoukry, M., Afzal, V., et al. (2013). Rapid and pervasive changes in genome-wide enhancer usage during mammalian development. Cell 155, 1521–1531. 10.1016/j.cell.2013.11.033

25. GTEx Consortium (2020). The GTEx Consortium atlas of genetic regulatory effects across human tissues. Science 369, 1318–1330. 10.1126/science.aaz1776

26. Schüttler, D., Bapat, A., Kääb, S., Lee, K., Tomsits, P., Clauss, S., and Hucker, W.J. (2020). Animal models of atrial fibrillation. Circ. Res. 127, 91–110. 10.1161/CIRCRESAHA.120.316366

27. Lang, D., Sulkin, M., Lou, Q., Efimov, I.R. (2011). Optical mapping of action potentials and calcium transients in the mouse heart. Journal of Visualized Experiments 55, 3275. 10.3791/3275

28. Lang, D., and Glukhov, A.V. (2016). High-resolution optical mapping of the mouse sino-atrial node. Journal of Visualized Experiments 118, 54773. 10.3791/54773

29. Grimes, K.M., Prasad, V., McNamara, J.W., Greaser, M.L., and Janssen, P.M.L. (2023). Rpl3l gene deletion in mice reduces heart weight over time. Front. Physiol. 14, 1054169. 10.3389/fphys.2023.1054169

30. Milenkoviæ, I., Zhang, Y., Drexler, H.C.A., Henning, R.H., and Krijgsveld, J. (2023). Dynamic interplay between RPL3- and RPL3L-containing ribosomes modulates mitochondrial activity in the mammalian heart. Nucleic Acids Res. 51, 5301–5324. 10.1093/nar/gkad121

31. Shiraishi, C., Matsumoto, A., Ichihara, K., Yamamoto, T., Yokoyama, T., Mizoo, T., Hatano, A., Matsumoto, M., Tanaka, Y., Matsuura-Suzuki, E., et al. (2023). RPL3L-containing ribosomes determine translation elongation dynamics required for cardiac function. Nat. Commun. 14, 2131. 10.1038/s41467-023-37838-6

32. Kao, B.R., Malerba, A., Lu-Nguyen, N.B., Harish, P., McCarthy, J.J., Dickson, G., and Popplewell, L.J. (2021). Knockdown of muscle-specific ribosomal protein L3-like enhances muscle function in healthy and dystrophic mice. Nucleic Acid Ther. 31, 457–464. 10.1089/nat.2020.0928

33. Xiang, Z., Thompson, A.D., Brogan, J.T., Schulte, M.L., Melancon, B.J., Mi, D., Lewis, L.M., Zou, B., Yang, L., Morrison, R., et al. (2011). The discovery and characterization of ML218: A novel, centrally active T-type calcium channel inhibitor with robust effects in STN neurons and in a rodent model of Parkinson’s disease. ACS Chem. Neurosci. 2, 730–742. 10.1021/cn200090z

34. Mangoni, M.E., Traboulsie, A., Leoni, A.L., Couette, B., Marger, L., Le Quang, K., Kupfer, E., Cohen-Solal, A., Vilar, J., Shin, H.S., et al. (2006). Bradycardia and slowing of the atrioventricular conduction in mice lacking CaV3.1/α1G T-type calcium channels. Circ. Res. 98, 1422–1430. 10.1161/01.RES.0000225862.14314.49

35. Harraz, O.F., Brett, S.E., Zechariah, A., Romero, M., Puglisi, J.L., Wilson, S.M., and Welsh, D.G. (2015). Genetic ablation of CaV3.2 channels enhances the arterial myogenic response by modulating the RyR-BKCa axis. Arterioscler. Thromb. Vasc. Biol. 35, 1843–1851. 10.1161/ATVBAHA.115.305736

36. Gupta, V., and Warner, J.R. (2014). Ribosome-omics of the human ribosome. RNA 20, 1004–1013. 10.1261/rna.043653.113

37. Chaillou, T., Zhang, X., and McCarthy, J.J. (2016). Expression of muscle-specific ribosomal protein L3-like impairs myotube growth. J. Cell. Physiol. 231, 1894–1902. 10.1002/jcp.25294

38. Hanssen, L.L.P., Kassouf, M.T., Oudelaar, A.M., Biggs, D., Preece, C., Downes, D.J., Gosden, M., Sharpe, J.A., Sloane-Stanley, J.A., Hughes, J.R., et al. (2017). Tissue-specific CTCF–cohesin-mediated chromatin architecture delimits enhancer interactions and function in vivo. Nat. Cell Biol. 19, 952–961. 10.1038/ncb3573

39. Barrington, C., Georgopoulou, D., Pezic, D., Varsally, W., Herrero, J., and Hadjur, S. (2019). Enhancer accessibility and CTCF occupancy underlie asymmetric TAD architecture and cell type specific genome topology. Nat. Commun. 10, 2908. 10.1038/s41467-019-10725-9

40. Vanden Broeck, A., and Klinge, S. (2024). Eukaryotic ribosome assembly. Annu. Rev. Biochem. 93, 189–210. 10.1146/annurev-biochem-030222-113611

41. Kim, T.-S., Kim, H.D., and Kim, J. (2009). PKCä-dependent functional switch of rpS3 between translation and DNA repair. Biochim. Biophys. Acta 1793, 395–405. 10.1016/j.bbamcr.2008.10.017

42. Park, Y.J., Kim, S.H., Kim, T.S., Lee, S.M., Cho, B.S., Seo, C.I., Kim, H.D., and Kim, J. (2021). Ribosomal protein S3 associates with the TFIIH complex and positively regulates nucleotide excision repair. Cell. Mol. Life Sci. 78, 3591–3606. 10.1007/s00018-020-03754-x

43. Mahata, B., Sundqvist, A., and Xirodimas, D.P. (2012). Recruitment of RPL11 at promoter sites of p53-regulated genes upon nucleolar stress through NEDD8 and in an Mdm2-dependent manner. Oncogene 31, 3060–3071. 10.1038/onc.2011.481

44. Nishimura, K., Kumazawa, T., Kuroda, T., Katagiri, N., Tsuchiya, M., Goto, N., Furumai, R., Murayama, A., Yanagisawa, J., and Kimura, K. (2015). Perturbation of ribosome biogenesis drives cells into senescence through 5S RNP-mediated p53 activation. Cell Rep. 10, 1310–1323. 10.1016/j.celrep.2015.01.055

45. Zhang, Y., O’Leary, M.N., Peri, S., Wang, M., Zha, J., Melov, S., Kappes, D.J., Feng, Q., Rhodes, J., Amieux, P.S., et al. (2017). Ribosomal proteins Rpl22 and Rpl22l1 control morphogenesis by regulating pre-mRNA splicing. Cell Rep. 18, 545–556. 10.1016/j.celrep.2016.12.034

46. Niwa, N., Yasui, K., Opthof, T., Takemura, H., Shimizu, A., Horiba, M., Lee, J.-K., Honjo, H., Kamiya, K., and Kodama, I. (2004). CaV3.2 subunit underlies the functional T-type Ca2+ channel in murine hearts during the embryonic period. Am. J. Physiol. Heart Circ. Physiol. 286, H2257–H2263. 10.1152/ajpheart.01043.2003

47. Yasui, K., Niwa, N., Takemura, H., Opthof, T., Muto, T., Horiba, M., Shimizu, A., Lee, J.-K., Honjo, H., Kamiya, K., et al. (2005). Pathophysiological significance of T-type Ca2+ channels: expression of T-type Ca2+ channels in fetal and diseased heart. J. Pharmacol. Sci. 99, 205–210. 10.1254/jphs.FMJ05002X3

48. Willemin, A., Szabo, D., and Pombo, A. (2024). Epigenetic regulatory layers in the 3D nucleus. Mol. Cell 84, 415–428. 10.1016/j.molcel.2023.12.003

49. Misteli, T. (2020). The self-organizing genome: principles of genome architecture and function. Cell 183, 28–45. 10.1016/j.cell.2020.09.014

50. Bersaglieri, C., and Santoro, R. (2019). Genome organization in and around the nucleolus. Cells 8, 579. 10.3390/cells8060579

51. Gupta, S., Goudarzi, K.M., Lin, J., Dieci, G., Babu, M.M., and Narlikar, G.J. (2025). The nucleolar granular component mediates genome organization through NPM1. Mol. Cell 85, 2782–2797.e8. 10.1016/j.molcel.2025.06.004

52. Xie, S.Q., Zhang, Y., So, C., Kim, E., Nam, Y., Jeong, J., and Kingston, R.E. (2022). Nucleolar-based Dux repression is essential for embryonic two-cell stage exit. Genes Dev. 36, 331–347. 10.1101/gad.349172.121

53. Al-Hassnan, Z.N., Almesned, A., Tulbah, S., Alakhfash, A., Alhadeq, F., Alruwaili, N., Alkorashy, M., Alhashem, A., Alrashdan, A., Faqeih, E., et al. (2020). Categorized genetic analysis in childhood-onset cardiomyopathy. Circ. Genom. Precis. Med. 13, 504–514. 10.1161/CIRCGEN.120.002969

54. Nannapaneni, H., Ghaleb, S., Arya, S., Gajula, V., Taylor, M.B., and Das, B.B. (2022). Further evidence of autosomal recessive inheritance of RPL3L pathogenic variants with rapidly progressive neonatal dilated cardiomyopathy. J. Cardiovasc. Dev. Dis. 9, 65. 10.3390/jcdd9030065

55. Das, B.B., Gajula, V., Arya, S., and Taylor, M.B. (2022). Compound heterozygous missense variants in RPL3L genes associated with severe forms of dilated cardiomyopathy: A case report and literature review. Children 9, 1495. 10.3390/children9101495

56. Yang, Q., Zhang, Q., Qin, Z., Zhang, S., Yi, S., Yi, S., Zhang, Q., and Luo, J. (2023). Novel compound heterozygous variants in the RPL3L gene causing dilated cardiomyopathy type-2D: A case report and literature review. BMC Med. Genomics 16, 127. 10.1186/s12920-023-01567-y

57. Bajpai, A.K., Gu, Q., Orgil, B.-O., Alberson, N.R., Towbin, J.A., Martinez, H.R., Lu, L., and Purevjav, E. (2023). Exploring the regulation and function of Rpl3l in the development of early-onset dilated cardiomyopathy and congestive heart failure using systems genetics approach. Genes 15, 53. 10.3390/genes15010053

58. Zhang, X., Wen, T., Chen, H., Jiang, Z., Gu, W., Yuan, W., Li, F., Shi, S., Shu, Q., and Yu, L. (2025). Novel compound heterozygous missense variants in RPL3L gene associated with neonatal dilated cardiomyopathy. Am. J. Med. Genet. A e64225. 10.1002/ajmg.a.64225

59. Murphy, M.R., Ganapathi, M., Lee, T.M., Fisher, J.M., Patel, M.V., Jayakar, P., Buchanan, A., Rippert, A.L., Ahrens-Nicklas, R.C., Nair, D., et al. (2025). Recessive but damaging alleles of muscle-specific ribosomes in dilated cardiomyopathy. bioRxiv. 10.1101/2025.01.02.630345

60. Agah, R., Frenkel, P.A., French, B.A., Michael, L.H., Overbeek, P.A., and Schneider, M.D. (1997). Gene recombination in postmitotic cells. Targeted expression of Cre recombinase provokes cardiac-restricted, site-specific rearrangement in adult ventricular muscle in vivo. J. Clin. Invest. 100, 169–179. 10.1172/JCI119509

61. Quignodon, L., Vincent, S., Winter, H., Samarut, J., and Flamant, F. (2007). A point mutation in the activation function 2 domain of thyroid hormone receptor á1 expressed after CRE-mediated recombination partially recapitulates hypothyroidism. Mol. Endocrinol. 21, 2350–2360. 10.1210/me.2007-0176

62. Ruan, H., Mandla, R., Ravi, N., Galang, G., Soe, A.W., Olgin, J.E., Lang, D., and Vedantham, V. (2023). Cholecystokinin-A signaling regulates automaticity of pacemaker cardiomyocytes. Front. Physiol. 14, 1284673. 10.3389/fphys.2023.1284673

63. Kleinsorge, M., and Cyganek, L. (2020). Subtype-directed differentiation of human iPSCs into atrial and ventricular cardiomyocytes. STAR Protoc. 1, 100026. 10.1016/j.xpro.2020.100026

64. Parasar, B., Raja Venkatesh, A., Perera, J., Sosnick, L., Moghadami, S., Seo, Y., Shi, J., Chan, L., Takenawa, S., Akiyama, T., et al. (2026). Whole-genome 3D architectural screen reveals modulators of brain DNA structure. bioRxiv. 10.64898/2026.04.15.718501

65. Luo, J., Deng, Z.-L., Luo, X., Tang, N., Song, W.-X., Chen, J., Sharff, K.A., Luu, H.H., Haydon, R.C., Kinzler, K.W., et al. (2007). A protocol for rapid generation of recombinant adenoviruses using the AdEasy system. Nat. Protoc. 2, 1236–1247. 10.1038/nprot.2007.135

